# Influence of Various Temporal Recoding on Pavlovian Eyeblink Conditioning in The Cerebellum

**DOI:** 10.1101/2020.06.23.168294

**Authors:** Sang-Yoon Kim, Woochang Lim

**Affiliations:** Institute for Computational Neuroscience and Department of Science Education, Daegu National University of Education, Daegu 42411, Korea

**Author notes:** Electronic address (Sang-Yoon Kim). Electronic address.

**Keywords:** Eyeblink conditioning, Effective learning, Various temporal recoding, Synaptic plasticity

## Abstract

We consider the Pavlovian eyeblink conditioning (EBC) via repeated presentation of paired conditioned stimulus (tone) and unconditioned stimulus (airpuff). The influence of various temporal recoding of granule cells on the EBC is investigated in a cerebellar network where the connection probability *p*_*c*_ from Golgi to granule cells is changed. In an optimal case of 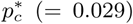, individual granule cells show various well- and ill-matched firing patterns relative to the unconditioned stimulus. Then, these variously-recoded signals are fed into the Purkinje cells (PCs) through parallel-fibers (PFs), and the instructor climbing-fiber (CF) signals from the inferior olive depress them effectively. In the case of well-matched PF-PC synapses, their synaptic weights are strongly depressed through strong long-term depression (LTD). On the other hand, practically no LTD occurs for the ill-matched PF-PC synapses. This type of “effective” depression at the PF-PC synapses coordinates firings of PCs effectively, which then make effective inhibitory coordination on cerebellar nucleus neuron [which elicits conditioned response (CR; eyeblink)]. When the learning trial passes a threshold, acquisition of CR begins. In this case, the timing degree 𝒯_*d*_ of CR becomes good due to presence of the ill-matched firing group which plays a role of protection barrier for the timing. With further increase in the trial, strength 𝒮 of CR (corresponding to the amplitude of eyelid closure) increases due to strong LTD in the well-matched firing group, while its timing degree 𝒯_*d*_ decreases. In this way, the well- and the ill-matched firing groups play their own roles for the strength and the timing of CR, respectively. Thus, with increasing the learning trial, the (overall) learning efficiency degree ℒ_*e*_ (taking into consideration both timing and strength of CR) for the CR is increased, and eventually it becomes saturated. By changing *p*_*c*_ from 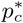, we also investigate the influence of various temporal recoding on the EBC. It is thus found that, the more various in temporal recoding, the more effective in learning for the Pavlovian EBC.

## I. INTRODUCTION

The cerebellum plays a crucial role in precise temporal and spatial motor control for coordination of voluntary movements (e.g., locomotion, balance, and posture), leading to smooth and balanced body movement [1–3]. In addition, it also participate in higher cognitive functions (e.g., attention, language, and speech) [2, 3]. The purpose of cerebellar motor learning is to carry out precise timing (associated with temporal information of movement) and gain (related to spatial information of movement) control for movements [4]. Experimental studies on timing and gain control for eye movements have been done in the two kinds of paradigms; (1) timing control for the Pavlovian eyeblink conditioning (EBC) [5–11] and (2) gain control for the vestibulo-ocular reflex and the optokinetic response [1, 12].

Here, we are interested in the Pavlovian EBC [see Fig. 1(a)] which is a representative example for the classical conditioning [13]. An animal (e.g., rabbit, rat, or mouse) receives repeated presentations of paired conditioned stimulus (CS; tone) and (eyeblink-eliciting) unconditioned stimulus (US; airpuff). When the training trial passes a threshold, the animal acquires the ability to elicit eyelid conditioned response (CR; acquisition of learned eyeblink) via learning representation of the time passage between the onsets of CS and US (i.e., the animal becomes conditioned to closes its eye in response to the tone CS with a time delay equal to the inter-stimulus interval (ISI) between the CS and the US onsets). The CRs exhibit two distinct features: (1) gradual acquisition of CR (i.e., CRs are acquired gradually over many training trials of repeated CS-US pairings) [14–18] and (2) adaptive timing of CR (i.e., CRs are well timed such that the time of peak eyelid closure matches well the ISI between the onsets of CS and US) [19–23]. Experimental works on EBC have been done in several animal species such as humans [24], monkeys [15], dogs [14], ferrets [18], rabbits [16], rats [17], and mice [25]. Particularly, since a series of groundbreaking experiments in rabbits [26, 27], EBC in restrained rabbits has served as a good model for motor learning.

**FIG. 1:**
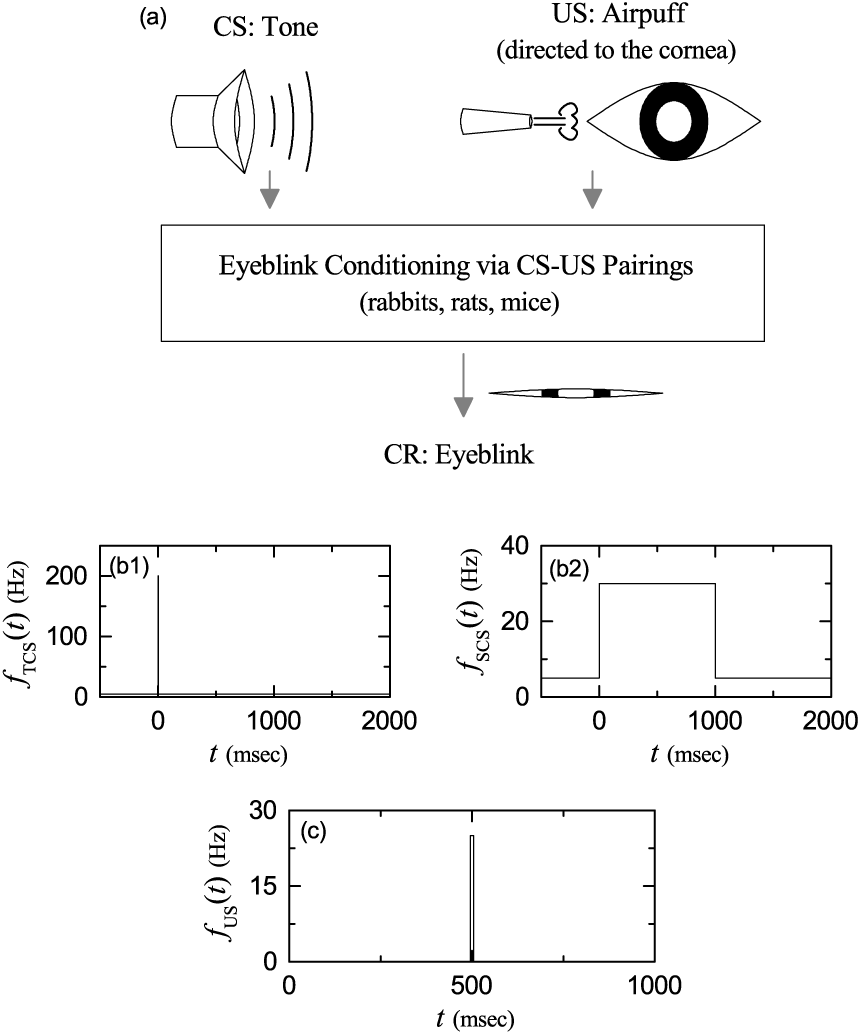
Pavlovian eyeblink conditioning (EBC). (a) Eyelid conditioned response (CR) (i.e., learned eyeblink) via repeated presentation of paired CS (conditioned stimulus) and US (unconditioned stimulus). Firing rates of (b1) transient conditioned stimulus (TCS) for resetting and (b2) sustained conditioned stimulus (SCS) (EBC signal). (c) Firing rate of transient unconditioned stimulus (US) for timing (eliciting unconditioned response).

Marr [28] and, later, Albus [29] formulated their seminal theory for cerebellar motor learning on the basis of its structure. Particularly, they paid attention to the recurrent network between the granule (GR) and the Golgi (GO) cells as a device of representation of spatial information (i.e., spatial coding). The input spatial patterns, conveyed via the mossy fibers (MFs), become more sparse and dissimilar to each other (i.e., pattern separation occurs) through recoding procedure in the granular layer composed of GR and GO cells. These recoded inputs are conveyed into the Purkinje cells (PCs) through the parallel fibers (PFs) (corresponding to the axons of GR cells). In addition to the PF recoded signals, the PCs also receive the error-teaching signals through the climbing-fiber (CF) from the inferior olive (IO). We assume that the PF-PC synapses are the only synapses where motor learning takes place. Thus, synaptic plasticity (i.e., potentiation or depression of synaptic strengths) may occur at PF-PC synapses. It is assumed by Marr [28] that a Hebbian type of long-term potentiation (LTP) occurs at the PF-PC synapses when both the PF and the CF signals are conjunctively excited [30, 31]. In opposition to Marr’s learning via LTP, it is assumed by Albus [29] that an anti-Hebbian type of long-term depression (LTD) takes place in the case of conjunctive excitations of both the PF and the CF signals. In the case of Albus’ learning via LTD, PCs learn when to stop their inhibition (i.e. when to disinhibit) rather than when to fire, because their firing activities become reduced. Several later experimental works have provided the support for the Albus’ learning via LTD [32–34]. Thus, LTD became established as a kind of synaptic plasticity for motor learning in the cerebellum [35–38].

A number of computational works for the EBC have been done. Several artificial models have been proposed for the time-passage representation (i.e., time coding) in the cerebellum [39–44]. However, these artificial models lacked biological plausibility. A realistic cerebellar model, based on many biological properties, has been built by focusing on the recurrent loop between the GR and the GO cells in the granular layer as a time-coding device [45]. Then, the realistic model generated a temporal code based on the population of active GR cells, and also, it was extended to reproduce the experimental results of the Pavlovian EBC [46, 47]. However, the computational mechanism to generate such a temporal code was unclear mainly due to complexity of the realistic model. To understand the computational mechanism for the time coding, a rate-coding model was developed in a simple recurrent inhibitory network, and its dynamics was analyzed in both the numerical and analytical way [48]. This rate-coding model generated a non-recurrent sequence of active neurons, corresponding to representation of a time-passage. Due to randomness in the recurrent connections, individual neurons exhibited random repetition of transitions between the active (bursting) and the inactive (silent) states which were persistent long-lasting ones. However, this rate-coding model is free of actual time scales. A spiking neural network model (with actual time scales) was built to examine representation of time passage in the cerebellum [49], and a large-scale computational simulation was also performed to reproduce some features of the EBC in the experiments.

However, the influence of various temporal recoding of GR cells on the Pavlovian EBC remains to be necessary for more clarification in several dynamical aspects, although much understanding on the representation of time passage in the cerebellum was achieved in previous computational works. As in the case of spatial coding for the optokinetic response [50], variously-recoded PF signals are dynamically classified for clear understanding of their matching with the instructor CF signals. Then, the dynamical classification of various firing patterns of GR cells leads us to clearly understand synaptic plasticity at PF-PC synapses and the following learning procedure. Consequently, we expect understanding on the rate of acquisition and the timing and strength of CR to be so much enhanced.

To this purpose, we employ a cerebellar ring network for the Pavlovian EBC, the basic framework of which was developed in the case of optokinetic response [50]. In such a cerebellar ring network, we vary the connection probability *p*_*c*_ from GO to GR cells and make a dynamical classification of various firing patterns of GR cells. GR cells in the whole population are divided into GR clusters. These GR clusters show various well- and ill-matched firing patterns with respect to the US (i.e., air-puff unconditioned stimulus) signal. Each firing pattern may be characterized in terms of the matching index between the firing pattern and the US. For quantifying the degree of various temporal recoding of GR cells, we introduce the variety degree 𝒱, given by the relative standard deviation in the distribution of matching indices of the firing patterns in the GR clusters, similar to the case of spatial coding for the optokinetic response [50]. We pay main attention to an optimal case of 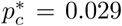 where the firing patterns of GR clusters are the most various. In the optimal case, 𝒱 ^*^ ≃ 1.842 which is a quantitative measure for various temporal recoding of GR cells. Dynamical origin of these various firing patterns of GR cells is also investigated. It is thus found that, various total synaptic inputs (including both the excitatory inputs via MFs and the inhibitory inputs from the pre-synaptic GO cells) into the GR clusters lead to generation of various firing patterns (i.e. outputs) in the GR clusters.

Based on dynamical classification of various firing patterns in the GR clusters, we employ a refined rule for synaptic plasticity (developed from the experimental result in [51]), and investigate intensively the influence of various temporal recoding of GR cells on synaptic plasticity at PF-PC synapses and subsequent learning process. PCs (corresponding to the cerebellar output) receive both the variously-recoded PF signals from the GR cells and the error-teaching CF signals from the IO neuron; the CF signals are well-matched with the US signal. In this case, CF and PF signals may be considered as “instructors” and “students,” respectively. Then, well-matched PF student signals are strongly depressed via strong LTD by the CF instructor signals. In contrast, practically no LTD occurs for the ill-matched PF student signals because most of them have no associations with the US signals which are strongly localized in the middle of each trial. In this way, the student PF signals are effectively depressed by the instructor CF signals.

During learning trials with repeated presentation of CS-US pairs, the “effective” depression at PF-PC synapses coordinates activations of PCs effectively, which then make effective inhibitory coordination on the cerebellar nucleus (CN) neuron [which elicits CR (i.e., learned eyeblink)]. In the optimal case of 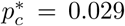, when passing the 141th threshold trial, acquisition of CR starts. In this optimal case, the timing degree 𝒯_*d*_ of the CR becomes good because of presence of ill-matched firing group which plays a role of protection barrier for the timing. With further increase in the trial, strength 𝒮 of CR [denoting the amplitude of eyelid closure (measured by the electromyography (EMG))] increases due to strong LTD in the well-matched firing group, while its timing degree 𝒯_*d*_ decreases. In this way, the well-and the ill-matched firing groups play their own roles for the strength and the timing of CR, respectively. Thus, the (overall) learning efficiency degree ℒ_*e*_ (considering both timing and strength of CR) for the CR increases with the training trial, and eventually it gets saturated.

Saturation in the learning procedure may be clearly shown in the inferior olive (IO) system. During the learning trials, the IO neuron receives both the excitatory US signal and the inhibitory input from the CN neuron. In this case, the learning progress degree ℒ_*p*_ is given by the ratio of the trial-averaged inhibitory input from the CN neuron to the magnitude of the trial-averaged excitatory input of the US signal. With increasing trial from the threshold, the trial-averaged inhibitory input (from the CN neuron) is increased, and approaches the constant trial-averaged excitatory input (through the US signal). Thus, at about the 250th trial, the learning progress degree becomes saturated at ℒ_*p*_ = 1. In this saturated case, the trial-averaged excitatory and inhibitory inputs to the IO neuron get balanced, and we obtain the saturated learning efficiency degree 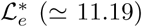 of the CR in the CN.

Finally, we vary *p*_*c*_ from 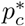, and investigate the influence of various temporal recoding of GR cells on the Pavlovian EBC. The distributions of both the variety degree 𝒱 of firing patterns of GR cells and the saturated learning efficiency degree 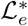 of CR versus *p*_*c*_ are found to form bell-shaped curves with peaks at 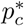. Moreover, 𝒱 both and 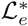 have a strong correlation. As a result, the more various in temporal recoding of GR cells, the more effective in learning for the Pavlovian EBC.

This paper is organized as follows. In Sec. II, a cerebellar ring network for the Pavlovian EBC is introduced. The governing equations for the population dynamics in the ring network are also given, together with a refined rule for the synaptic plasticity at the PF-PC synapses. Then, in the main Sec. III, we investigate the influence of various temporal recoding of GR cells on learning for the Pavlovian EBC by varying *p*_*c*_. Finally, summary and discussion are given in Sec. IV.

## II. CEREBELLAR RING NETWORK FOR THE PAVLOVIAN EYEBLINK CONDITIONING

In this section, we describe our cerebellar ring network for the Pavlovian EBC. The basic framework of such a ring network was developed in the case of optokinetic response [50]. The ring network for the EBC is essentially the same as that for the optokinetic response, except for the number of glomeruli in each GR cluster which is 4 and 2 for the EBC and the optokinetic response, respectively. Also, the external input signals (i.e., the context MF signal and the desired IO signal) for the EBC are different from those for the optokinetic response. For the sake of completeness, we also include a detailed explanation on the cerebellar ring network within this paper.

### A. Conditioned Stimulus and Unconditioned Stimulus

Figure 1(a) shows the Pavlovian EBC. During the training trials, repeated presentations of paired tone CS and delayed airpuff US are made to an animal (e.g., rabbit, rat, or mouse). As the training trial passes a threshold, the animal acquires the ability to elicit eyelid CR (i.e., acquisition of learned eyeblink) through learning representation of the time passage between the CS and the US onsets. Accordingly, the animal gets conditioned to closes its eye in response to the tone CS with a time delay equal to the ISI between the onsets of CS and US.

In this subsection, we give explanations on the two external input signals. We first consider the CS for the EBC signal. When the CS is a tone, the pontine nucleus in the pons receives the auditory information, and then it sends the “context” signal for the EBC via MFs to both the GR cells and the CN neuron. There are a transient CS for resetting and a sustained CS (representing the tone) [49]. Each step (0 < *t* < 2000 msec) for EBC learning consists of the trial stage (0 < *t* < 1000 msec) and the break stage (1000 < *t* < 2000 msec); *t* denotes the time. In the trial stage, the transient CS is modeled in terms of strong and brief Poisson spike trains of 200 Hz for 0 < *t* < 5 msec and the subsequent background Poisson spike trains of 5 Hz for 5 < *t* < 1000 msec. On the other hand, the sustained CS is modeled in terms of Poisson spike trains of 30 Hz for 0 < *t* < 1000 msec. In the following break stage, the CS is modeled in terms of the background Poisson spike trains of 5 Hz for 1000 < *t* < 2000 msec. The firing rates *f*_TCS_(*t*) and *f*_SCS_(*t*) of the transient CS and the sustained CS are shown in Figs. 1(b1) and 1(b2), respectively. These figures also show the preparatory step for − 500 < *t* < 0 msec where the CS is modeled in terms of the background Poisson spike trains of 5 Hz; this preparatory step precedes just the 1st step for the EBC learning.

Next, we consider the US for the desired timing signal. When an airpuff US is delivered to the cornea of an eye, sensory information is carried to the sensory trigeminal nucleus (which extends through the whole of midbrain, pons, and medulla and into the high cervical spinal cord). Then, the trigeminal nucleus also sends the desired timing signal to the IO. In the trial stage (0 < *t* < 1000 msec), the US (eliciting unconditioned response) is modeled in terms of the strong and brief Poisson spike trains of 25 Hz for a short interval in the middle of the trial stage, *t*^*^− Δ*t* < *t* < *t*^*^ + Δ*t* (*t*^*^ = 500 msec and Δ*t* = 5 msec) [49]. The firing rate *f*_US_(*t*) of the US is shown in Fig. 1(c).

### B. Framework for Cerebellar Ring Network for The Pavlovian EBC

A cerebellar ring network was introduced in the case of optokinetic response [50]. As in the famous small-world ring network [52, 53], a one-dimensional ring network with a simple architecture was developed, which is in contrast to the two-dimensional square-lattice network [4, 49]. This kind of ring network has advantage for computational and analytical efficiency, and its visual representation may also be easily made.

Here, we employ such a cerebellar ring network for the Pavlovian EBC. Figure 2(a) shows the box diagram for the cerebellar network. The granular layer, corresponding to the input layer of the cerebellar cortex, is composed of the excitatory GR cells and the inhibitory GO cells. On the other hand, the Purkinje-molecular layer, corresponding to the output layer of the cerebellar cortex, consist of the inhibitory PCs and the inhibitory BCs (basket cells). The MF context signals for the EBC are fed from the pontine nucleus in the pons to the GR cells; each GR cell receives two transient and two sustained CS signals via four MFs (i.e., two pairs of transient and sustained CS signals are fed into each GR cell). Various temporal recoding is made in the granular layer via inhibitory coordination of GO cells on GR cells. Then, these various-recoded outputs are fed via PFs to the PCs and the BCs in the Purkinje-molecular layer.

**FIG. 2:**
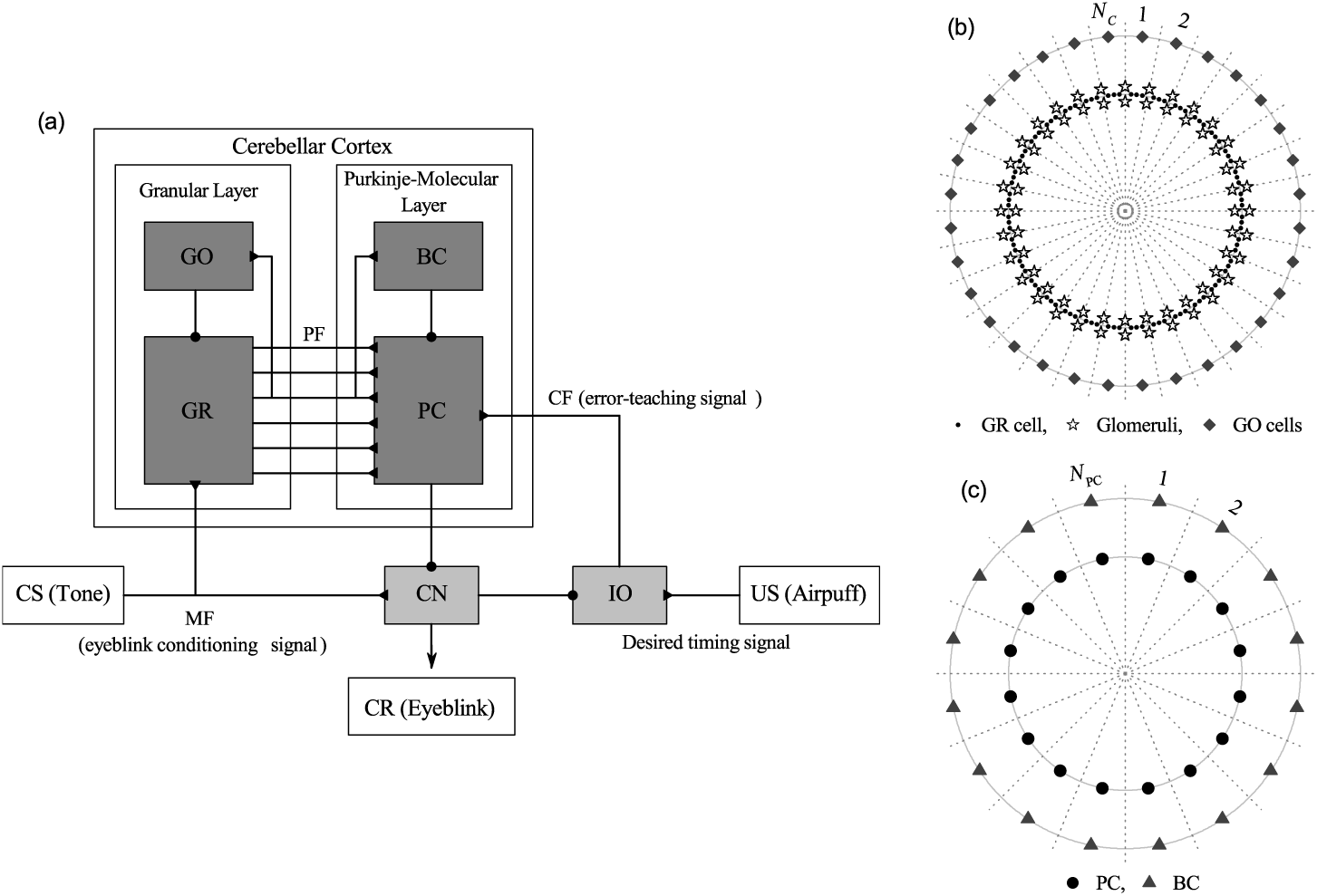
Cerebellar ring network for the EBC. (a) Box diagram for the cerebellar network. Lines with triangles and circles represent excitatory and inhibitory synapses, respectively. GR (granule cell), GO (Golgi cell), and PF (parallel fiber) in the granular layer, PC (Purkinje cell) and BC (basket cell) in the Purkinje-molecular layer, and other parts for CN (cerebellar nuclei), IO(inferior olive), MF (mossy fiber), and CF (climbing fiber). (b) Schematic diagram for granular-layer ring network with concentric inner GR and outer GO rings. Numbers represent granular layer zones (bounded by dotted lines) for *N*_*C*_ = 32. In each *I*th zone (*I* = 1,, *N*_*C*_), there exists the *I*th GR cluster on the inner GR ring. Each GR cluster consists of GR cells (solid circles), and it is bounded by 4 glomeruli (stars). On the outer GO ring in the *I*th zone, there exists the *I*th GO cell (diamonds). (c) Schematic diagram for Purkinje-molecular-layer ring network with concentric inner PC and outer BC rings. Numbers represent the Purkinje-molecular-layer zones (bounded by dotted lines) for *N*_PC_ = 16. In each *J* th zone, there exist the *J* th PC (solid circle) on the inner PC ring and the *J* th BC (solid triangle) on the outer BC ring.

The PCs receive another excitatory error-teaching CF signals from the IO, along with the inhibitory inputs from the BCs. Then, depending on the type of PF signals (i.e., well- or ill-matched PF signals), various PF (student) signals are effectively depressed by the error-teaching (instructor) CF signals. Such “effective” depression at the PF-PC synapses coordinates firings of PCs effectively, which then exert effective inhibitory coordination on the CN neuron. The CN neuron also receives two excitatory signals; one transient and one sustained CS signals via MFs. In the earlier trials, the CN neuron can not fire, due to strong inhibition from the PCs. As the learning trial passes a threshold, the CN neuron starts firing, and then it exerts excitatory projections onto the eyeblink pre-motoneurons in the midbrain which then supply motor commands to eyeblink motoneurons. Thus, acquisition of CR begins (i.e., acquisition of learned eyeblink starts). This CN neuron also provides inhibitory inputs to the IO neuron which also receives the excitatory signals for the desired timing from the trigeminal nucleus. Then, the IO neuron supplies excitatory error-teaching CF signals to the PCs.

Figure 2(b) shows a schematic diagram for the granular-layer ring network with concentric inner GR and outer GO rings. Numbers represent granular-layer zones (bounded by dotted lines); the numbers 1, 2, *…*, and *N*_*C*_ represent the 1st, the 2nd, *…*, and the *N*_*C*_ th granular-layer zones, respectively. Thus, the total number of granular-layer zones is *N*_*C*_ ; Fig. 2(b) shows an example for *N*_*C*_ = 32. In each *I*th zone (*I* = 1, *…, N*_*C*_), there exists the *I*th GR cluster on the inner GR ring. Each GR cluster consists of *N*_GR_ excitatory GR cells (solid circles). Then, location of each GR cell may be denoted by the two indices (*I, i*) which represent the *i*th GR cell in the *I*th GR cluster, where *i* = 1, *…, N*_GR_. Here, we consider the case of *N*_*C*_ = 2^10^ and *N*_GR_ = 50, and thus the total number of GR cells is 51,200. In this case, the *I*th zone covers the angular range of 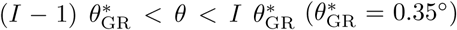. On the outer GO ring in each *I*th zone, there exists the *I*th inhibitory GO cell (diamond), and thus the total number of GO cells is *N*_*C*_.

We note that each GR cluster is bounded by 4 glomeruli (corresponding to the axon terminals of the MFs) (stars) at both boundaries of the GR cluster; at each boundary, a pair of glomeruli (upper and lower ones) exist. This is in contrast to the case of optokinetic response where 2 glomeruli bound each GR cluster; at each left or right boundary, only one glomerulus is located. GR cells within each GR cluster share the same inhibitory and excitatory synaptic inputs through their dendrites which contact the four glomeruli at both ends of the GR cluster. Each glomerulus receives inhibitory inputs from nearby 81 (clockwise side: 41 and counter-clockwise side: 40) GO cells with a random connection probability *p*_*c*_ (= 0.029). Hence, on average, about 2 GO cell axons innervate each glomerulus. Thus, each GR cell receives about 9 inhibitory inputs through 4 dendrites which synaptically contact the glomeruli at both boundaries. In this way, each GR cell in the GR cluster shares the same inhibitory synaptic inputs from nearby GO cells through the intermediate glomeruli at both ends.

In addition, each GR cell shares the same four excitatory inputs via the four glomeruli at both boundaries, because a glomerulus receives an excitatory MF input. We note that transient CS signals are supplied via the two upper glomeruli, while sustained CS signals are fed through the two lower glomeruli. Here, we take into consideration stochastic variability of synaptic transmission from a glomerulus to GR cells, and supply independent Poisson spike trains with the same firing rate to each GR cell for the excitatory MF signals. In this GR-GO feedback system, each GO cell receives excitatory synaptic inputs through PFs from GR cells in the nearby 49 (central side: 1, clockwise side: 24 and counter-clockwise side: 24) GR clusters with a random connection probability 0.1. Hence, 245 PFs (i.e. GR cell axons) innervate a GO cell.

Figure 2(c) shows a schematic diagram for the Purkinje-molecular-layer ring network with concentric inner PC and outer BC rings. Numbers represent the Purkinje-molecular-layer zones (bounded by dotted lines). In each *J* th zone (*J* = 1, *…, N*_PC_), there exist the *J* th PC (solid circles) on the inner PC ring and the *J* th BC (solid triangles) on the outer BC ring. Here, we consider the case of *N*_PC_ = 16, and thus the total numbers of PC and BC are 16, respectively. In this case, each *J* th (*J* = 1, *…, N*_PC_) zone covers the angular range of (*J* - 1) 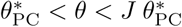, where 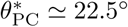 (corresponding to about 64 zones in the granular-layer ring network). We note that variously-recoded PFs innervate PCs and BCs. Each PC (BC) in the *J* th Purkinje-molecular-layer zone receives excitatory synaptic inputs via PFs from all the GR cells in the 288 GR clusters (clockwise side: 144 and counter-clockwise side: 144 when starting from the angle 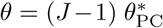 in the granular-layer ring network). Thus, each PC (BC) is synaptically connected via PFs to the 14,400 GR cells (which corresponds to about 28 % of the total GR cells). In addition to the PF signals, each PC also receives inhibitory inputs from nearby 3 BCs (central side: 1, clockwise side: 1 and counter-clockwise side: 1) and excitatory error-teaching CF signal from the IO.

Here, for simplicity, we consider just one CN neuron and one IO neuron. Both excitatory inputs (corresponding to one transient and one sustained CS signals) via 2 MFs and inhibitory inputs from all the 16 PCs are fed into the CN neuron. Then, the CN neuron provides excitatory input to the eyeblink pre-motoneurons in the mid-brain and also supplies inhibitory input to the IO neuron. One additional excitatory desired timing signal from the trigeminal nucleus is also fed into the IO neuron. Then, through integration of both excitatory and inhibitory inputs, the IO neuron provides excitatory error-teaching CF signals to the PCs.

### C. Elements of The Cerebellar Ring Network

As in the case of optokinetic response [50], we choose leaky integrate-and-fire (LIF) neuron models as elements of the cerebellar ring network [54]. Here, the LIF neuron models incorporate additional afterhyperpolarization (AHP) currents that determine refractory periods. This LIF neuron model is one of the simplest spiking neuron models. Because of its simplicity, it may be easily analyzed and simulated. Hence, it has been very popularly employed as a neuron model.

Dynamics of states of individual neurons in the *X* population are governed by the following equations:

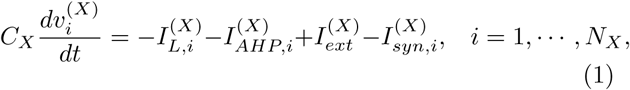

where *N*_*X*_ is the total number of neurons in the *X* population, *X* = GR and GO in the granular layer, *X* = PC and BC in the Purkinje-molecular layer, and in the other parts *X* = CN and IO. The state of the *i*th neuron in the *X* population at a time *t* (msec) is characterized by its membrane potential 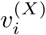 (mV), and *C*_*X*_ (pF) denotes the membrane capacitance of the cells in the *X* population. The time-evolution of 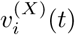 is governed by 4 types of currents (pA) into the *i*th neuron in the *X* population; the leakage current 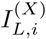, the AHP current 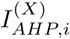, the external constant current 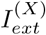 (independent of *i*), and the synaptic current 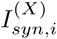.

We consider a single LIF neuron model [without the AHP current and the synaptic current in Eq. (1)] which describes a simple parallel resistor-capacitor circuit. Here, the leakage term is due to the resistor and the integration of the external current is due to the capacitor which is in parallel to the resistor. Thus, in Eq. (1), the 1st type of leakage current 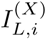 for the *i*th neuron in the *X* population is given by:

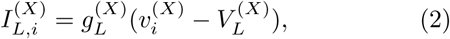

where 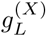 and 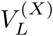 are conductance (nS) and reversal potential for the leakage current, respectively.

The *i*th neuron fires a spike when its membrane potential 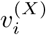 reaches a threshold 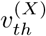 at a time 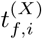. Then,the 2nd type of AHP current 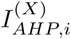 follows after firing (i.e., 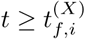):

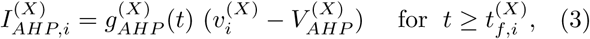

where 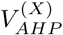 AHP is the reversal potential for the AHP current. The conductance 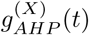 is given by an exponential-decay function:

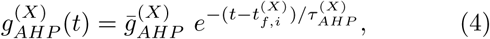

where 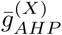 and 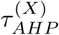 are the maximum conductance and the decay time constant for the AHP current. As 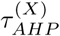 increases, the refractory period becomes longer.

The 3rd type of external constant current 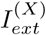 for spontaneous firing is provided to only PCs because of their high spontaneous firing rate [55, 56]. In Appendix, Table I shows the parameter values for the capacitance *C*_*X*_, the leakage current 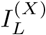, the AHP current 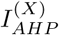, and the external constant current 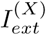. These values are adopted from physiological data [49].

**TABLE I:**
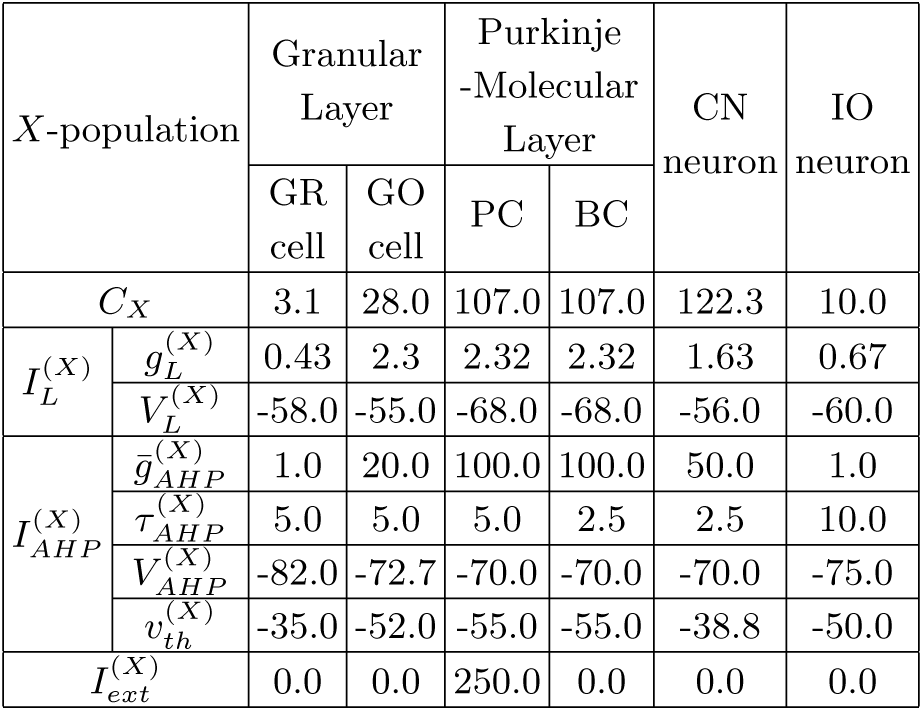
Parameter values for LIF neuron models with AHP currents for the granule (GR) cell and the Golgi (GO) cell in the granular layer, the Purkinje cell (PC) and the basket cell (BC) in the Purkinje-molecular layer, and the cerebellar nucleus (CN) and the inferior olive (IO) neurons.

| X-population |  | Granular Layer |  | Purkinje -Molecular Layer |  | CN neuron | IO neuron |
| --- | --- | --- | --- | --- | --- | --- | --- |
|  |  | GR cell | GO cell | PC | BC |  |  |
| $C_X$ | | 3.1 | 28.0 | 107.0 | 107.0 | 122.3 | 10.0 |
| $I_L^{(X)}$ | $g_L^{(X)}$ | 0.43 | 2.3 | 2.32 | 2.32 | 1.63 | 0.67 |
| | $V_L^{(X)}$ | -58.0 | -55.0 | -68.0 | -68.0 | -56.0 | -60.0 |
| $I_{AHP}^{(X)}$ | $\bar{g}_{AHP}^{(X)}$ | 1.0 | 20.0 | 100.0 | 100.0 | 50.0 | 1.0 |
| | $\tau_{AHP}^{(X)}$ | 5.0 | 5.0 | 5.0 | 2.5 | 2.5 | 10.0 |
| | $V_{AHP}^{(X)}$ | -82.0 | -72.7 | -70.0 | -70.0 | -70.0 | -75.0 |
| | $v_{th}^{(X)}$ | -35.0 | -52.0 | -55.0 | -55.0 | -38.8 | -50.0 |
| $I_{ext}^{(X)}$ | | 0.0 | 0.0 | 250.0 | 0.0 | 0.0 | 0.0 |

### D. Three Kinds of Synaptic Currents

Here, we are concerned about the 4th type of synaptic current 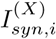 into the *i*th neuron in the *X* population in Eq. (1). It is composed of the following 3 kinds of synaptic currents:

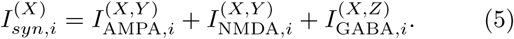

Here, 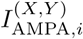 and 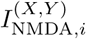 are the excitatory AMPA (*α*-amino-3-hydroxy-5-methyl-4-isoxazolepropionic acid) receptor-mediated and NMDA (*N* -methyl-*D*-aspartate) receptor-mediated currents from the pre-synaptic source *Y* population to the post-synaptic *i*th neuron in the target *X* population. In contrast, 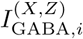 is the inhibitory GABA_A_ (*γ*-aminobutyric acid type A) receptor-mediated current from the pre-synaptic source *Z* population to the post-synaptic *i*th neuron in the target *X* population.

As in the case of the AHP current, the *R* (= AMPA, NMDA, or GABA) receptor-mediated synaptic current 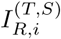 from the pre-synaptic source *S* population to the *i*th post-synaptic neuron in the target *T* population is given by:

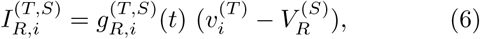

where 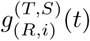 and 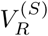 are synaptic conductance and synaptic reversal potential (determined by the type of the pre-synaptic source *S* population), respectively. We obtain the synaptic conductance 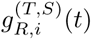 from:

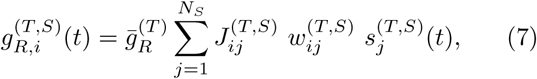

where 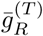 and 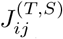 are the maximum conductance and the synaptic weight of the synapse from the *j*th pre-synaptic neuron in the source *S* population to the *i*th post-synaptic neuron in the target *T* population, respectively. The inter-population synaptic connection from the source *S* population (with *N*_*s*_ neurons) to the target *T* population is given in terms of the connection weight matrix 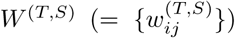 where 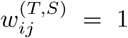 if the *j*th neuron in the source *S* population is pre-synaptic to the *i*th neuron in the target *T* population; otherwise 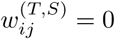.

The post-synaptic ion channels are opened because of the binding of neurotransmitters (emitted from the source *S* population) to receptors in the target *T* population. The fraction of open ion channels at time *t* is represented by *s*^(*T,S*)^. The time course of 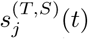 of the *j*th neuron in the source *S* population is given by a sum of exponential-decay functions 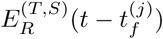:

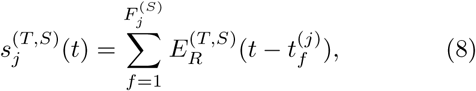

where 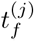 and 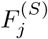 are the *f* th spike time and the total number of spikes of the *j*th neuron in the source *S* population, respectively. The exponential-decay function 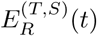 (which corresponds to contribution of a pre-synaptic spike occurring at *t* = 0 in the absence of synaptic delay) is given by:

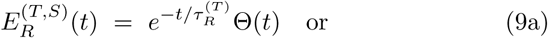

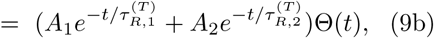

where Θ(*t*) is the Heaviside step function: Θ(*t*) = 1 for *t* ≥ 0 and 0 for *t* < 0. Depending on the source and the target populations, 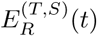 may be a type-1 single exponential-decay function of Eq. (9a) or a type-2 dual exponential-decay function of Eq. (9b). In the type-1 case, there exists one synaptic decay time constant 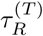 (determined by the receptor on the post-synaptic target *T* population), while in the type-2 case, two synaptic decay time constants, 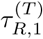 and 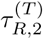 appear. In most cases, the type-1 single exponential-decay function of Eq. (9a) appears, except for the two synaptic currents 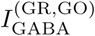 and 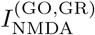.

In Appendix, Table II shows the parameter values for the maximum conductance 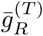, the synaptic weight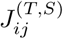, the synaptic reversal potential 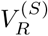, the synaptic decay time constant 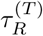, and the amplitudes *A*_1_ and *A*_2_ for the type-2 exponential-decay function in the granular layer, the Purkinje-molecular layer, and the other parts for the CN and IO, respectively. These values are adopted from physiological data [49].

**TABLE II:** Parameter values for synaptic currents 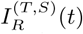 into the granule (GR) and the Golgi (GO) cells in the granular layer, the Purkinje cells (PCs) and the basket cells (BCs) in the Purkinje-molecular layer, and the cerebellar nucleus (CN) and the inferior olive (IO) neurons in the other parts. In the granular layer, the GR cells receive excitatory inputs via mossy fibers (MFs) and inhibitory inputs from GO cells, and the GO cells receive excitatory inputs via parallel fibers (PFs) from GR cells. In the Purkinje-molecular layer, the PCs receive two types of excitatory inputs via PFs from GR cells and through climbing fibers (CFs) from the IO and one type of inhibitory inputs from the BCs. The BCs receive excitatory inputs via PFs from GR cells. In the other parts, the CN neuron receives excitatory inputs via MFs and inhibitory inputs from PCs, and the IO neuron receives excitatory input via the US signal and inhibitory input from the CN neuron.

|  | Granular Layer |  |  |  |  | Purkinje-Molecular Layer |  |  |  | Other Parts |  |  |  |  |
| --- | --- | --- | --- | --- | --- | --- | --- | --- | --- | --- | --- | --- | --- | --- |
| Target Cells ($T$) | GR | | | GO | | PC | | | BC | CN | | | IO | |
| Source Cells ($S$) | MF | MF | GO | PF | PF | PF | CF | BC | PF | MF | MF | PC | US | CN |
| Receptor ($R$) | AMPA | NMDA | GABA | AMPA | NMDA | AMPA | AMPA | GABA | AMPA | AMPA | NMDA | GABA | AMPA | GABA |
| $\bar{g}_R^{(T)}$ | 0.18 | 0.025 | 0.028 | 45.5 | 30.0 | 0.7 | 0.7 | 1.0 | 0.7 | 50.0 | 25.8 | 30.0 | 1.0 | 0.18 |
| $J_{ij}^{(T,S)}$ | 8.0 | 8.0 | 10.0 | 0.00004 | 0.00004 | 0.006 | 1.0 | 5.3 | 0.006 | 0.002 | 0.002 | 0.008 | 1.0 | 5.0 |
| $V_R^{(S)}$ | 0.0 | 0.0 | -82.0 | 0.0 | 0.0 | 0.0 | 0.0 | -75.0 | 0.0 | 0.0 | 0.0 | -88.0 | 0.0 | -75.0 |
| $\tau_R^{(T)}$ | 1.2 | 52.0 | 7.0, 59.0 | 1.5 | 31.0, 170.0 | 8.3 | 8.3 | 10.0 | 8.3 | 9.9 | 30.6 | 42.3 | 10.0 | 10.0 |
| $A_1, A_2$ | | | 0.43, 0.57 | | 0.33, 0.67 | | | | | | | | | |

### E. Refined Rule for Synaptic Plasticity

As in [50], we employ a rule for synaptic plasticity, based on the experimental result in [51]. This rule is a refined one for the LTD in comparison with the rule used in [4, 49], the details of which will be explained below.

The coupling strength of the synapse from the pre-synaptic neuron *j* in the source *S* population to the post-synaptic neuron *i* in the target *T* population is 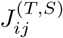. Initial synaptic strengths for 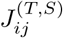 are given in Table II In this work, we assume that learning occurs only at the PF-PC synapses. Hence, only the synaptic strengths 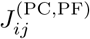 of PF-PC synapses may be modifiable (i.e., they are depressed or potentiated), while synaptic strengths of all the other synapses are static. [Here, the index *j* for the PFs corresponds to the two indices (*M, m*) for GR cells representing the *m*th (1 ≤*m* ≤ 50) cell in the *M* th (1 ≤ *M* ≤ 2^10^) GR cluster.] Synaptic plasticity at PF-PC synapses have been so much studied in many experimental [32–35, 51, 57–65] and computational [4, 29, 49, 66–72] works.

As the time *t* is increased, synaptic strength 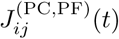 for each PF-PC synapse is updated with the following multiplicative rule (depending on states) [50, 51]:

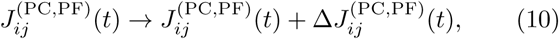

Where

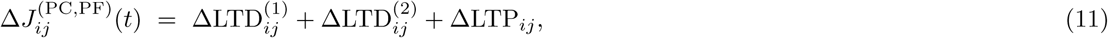

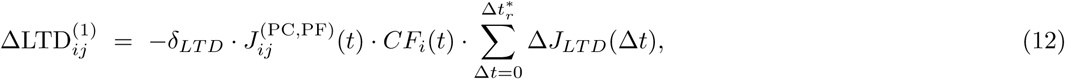

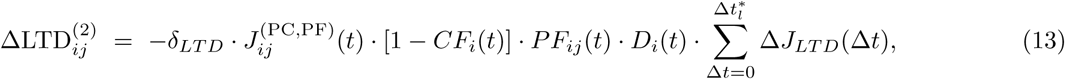

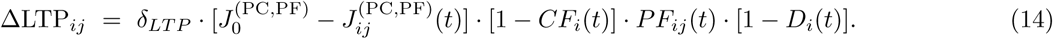

Here, 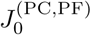 is the initial value (=0.006) for the synaptic strength of PF-PC synapses. Synaptic modification (LTD or LTP) occurs, depending on the relative time difference Δ*t* [= *t*_CF_ (CF activation time) - *t*_PF_ (PF activation time)] between the spiking times of the error-teaching instructor CF and the variously-recoded student PF. In Eqs. (12)-(14), *CF*_*i*_(*t*) denotes a spike train of the CF signal coming into the *i*th PC. When *CF*_*i*_(*t*) activates at a time *t, CF*_*i*_(*t*) = 1; otherwise, *CF*_*i*_(*t*) = 0. This instructor CF activation gives rise to LTD at PFPC synapses in conjunction with earlier (Δ*t* > 0) student PF activations in the range of 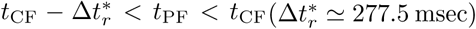, which corresponds to the major LTD in Eq. (12).

We next consider the case of *CF*_*i*_(*t*) = 0, which corresponds to Eqs. (13) and (14). Here, *PF*_*ij*_(*t*) denotes a spike train of the PF signal from the *j*th pre-synaptic GR cell to the *i*th post-synaptic PC. When *PF*_*ij*_(*t*) activates at time *t, PF*_*ij*_(*t*) = 1; otherwise, *PF*_*ij*_(*t*) = 0. In the case of *PF*_*ij*_(*t*) = 1, PF firing may cause LTD or LTP, depending on the presence of earlier CF firings in an effective range. If CF firings exist in the range of 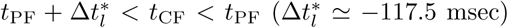, *D*_*i*_(*t*) = 1; otherwise *D*_*i*_(*t*) = 0. When both *PF*_*ij*_(*t*) = 1 and *D*_*i*_(*t*) = 1, the PF activation causes another LTD at PF-PC synapses in conjunction with earlier (Δ*t* < 0) CF activations [see Eq. (13)]. The probability for occurrence of earlier CF firings within the effective range is very low because mean firing rates of the CF signals (corresponding to output firings of individual IO neurons) are ∼ 1.5 Hz [73, 74]. Hence, this 2nd type of LTD is a minor one. In contrast, in the case of *D*_*i*_(*t*) = 0 (i.e., absence of earlier associated CF firings), LTP occurs because of the PF firing alone [see Eq. (14)]. The update rate *δ*_*LT D*_ for LTD in Eqs. (12) and (13) is 0.005, while the update rate *δ*_*LT P*_ for LTP in Eqs. (14) is 0.0005 (=*δ*_*LT D*_*/*10) [4].

In the case of LTD in Eqs. (12) and (13), the synaptic modification Δ*J*_*LT D*_(Δ*t*) changes depending on the relative time difference Δ*t* (= *t*_CF_ − *t*_PF_). We use the following time window for the synaptic modification Δ*J*_*LT D*_(Δ*t*) [50, 51]:

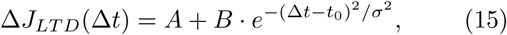

where *A* = −0.12, *B* = 0.4, *t*_0_ = 80, and *σ* = 180.

Figure 3 shows the time window for Δ*J*_*LTD*_(Δ*t*). As shown well in Fig. 3, LTD occurs in an effective range of 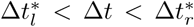. We note that a peak appears at *t*_0_ = 80 msec, and hence peak LTD takes place when PF firing precedes CF firing by 80 msec. A CF firing gives rise to LTD in association with earlier PF firings in the region hatched with horizontal lines 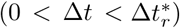, and it also causes to another LTD in conjunction with later PF firings in the region hatched with vertical lines 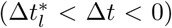. The effect of CF firing on earlier PF firings is much larger than that on later PF firings. However, outside the effective range (i.e., 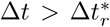 or 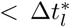), PF firings alone results in occurrence of LTP, because of absence of effectively associated CF firings.

**FIG. 3:**
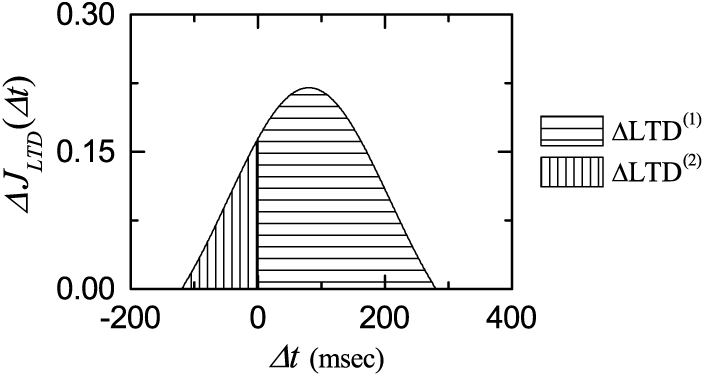
Time window for the LTD at the PF-PC synapse. Plot of synaptic modification Δ*J*_*LTD*_ (Δ*t*) for LTD versus Δ*t*.

Our refined rule for synaptic plasticity has the following advantages for the ΔLTD in comparison with that in [4, 49]. Our rule is based on the experimental result in [51]. In the presence of a CF firing, a major LTD (ΔLTD^(1)^) occurs in conjunction with earlier PF firings in the range of 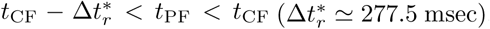, while a minor LTD (ΔLTD^(2)^) takes place in conjunction with later PF firings in the range of 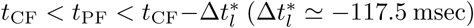. The magnitude of LTD varies depending on Δ*t* (= *t*_CF_ - *t*_PF_); a peak LTD occurs when Δ*t* = 80 msec. In contrast, the rule in [4, 49] considers only the major LTD in association with earlier PF firings in the range of *t*_CF_ − 50 < *t*_PF_ < *t*_CF_, the magnitude of major LTD is equal, independently of Δ*t*, and minor LTD in conjunction with later PF firings is not considered. Outside the effective range of LTD, PF firings alone lead to LTP in both rules.

### F. Numerical Method

Numerical integration of the governing Eq. (1) for the time-evolution of states of individual neurons, together with the update rule for synaptic plasticity of Eq. (10), is made by using the 2nd-order Runge-Kutta method with the time step 1 msec. In each realization, we choose random initial points 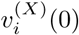 for the *i*th neuron in the *X* population with uniform probability in the range of 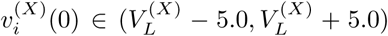 ; the values of 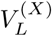 are given in Table I.

## III. INFLUENCE OF VARIOUS TEMPORAL RECODING IN GR CLUSTERS ON LEARNING FOR THE PAVLOVIAN EYEBLINK CONDITIONING

In this section, we investigate the influence of various temporal recoding of GR cells on learning for the EBC by changing the connection probability *p*_*c*_ from the GO to the GR cells. We mainly focus on an optimal case of 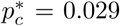 where the firing patterns of GR clusters are the most various. In this case, we first make dynamical classification of various firing patterns of the GR clusters. Next, we study the influence of various firing patterns on the synaptic plasticity at the PF-PC synapses and the subsequent learning process in the PC-CN-IO system. Finally, we change *p*_*c*_ from the optimal value 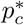, and investigate dependence of the variety degree 𝒱 of firing patterns and the saturated learning efficiency degree 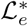 for the CR on *p*_*c*_. Both 𝒱 and 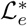 are found to form bell-shaped curves with peaks at 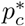, and they have strong correlation with the Pearson’s coefficient *r* ≃ 0.9982 [75]. Consequently, the more various in temporal recoding in the GR clusters, the more effective in the learning for the EBC.

### A. Collective Firing Activity in The Whole Population of GR Cells

Temporal recoding process is performed in the granular layer (corresponding to the input layer of the cerebellar cortex), composed of GR and GO cells (see Fig. 2). GR cells (principal output cells in the granular layer) receive excitatory context signals for the EBC via the MFs [see Figs. 1(b1) and 1(b2)] and make various recoding of context signals through receiving effective inhibitory coordination of GO cells. Thus, variously recoded signals are fed into the PCs (principal output cells in the cerebellar cortex) via PFs.

We first consider the firing activity in the whole population of GR cells for 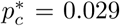. Collective firing activity may be well visualized in the raster plot of spikes which is a collection of spike trains of individual neurons. As a collective quantity showing whole-population firing behaviors, we use an instantaneous whole-population spike rate *R*_GR_(*t*) which may be got from the raster plots of spikes [76–83]. To get a smooth instantaneous whole-population spike rate, we employ the kernel density estimation (kernel smoother) [84]. Each spike in the raster plot is convoluted (or blurred) with a kernel function *K*_*h*_(*t*), and then a smooth estimate of instantaneous whole-population spike rate *R*_GR_(*t*) is got by averaging the convoluted kernel function over all spikes of GR cells in the whole population:

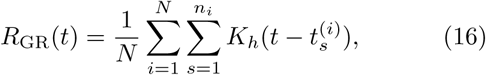

where 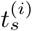 is the *s*th spiking time of the *i*th GR cell, *n*_*i*_ is the total number of spikes for the *i*th GR cell, and *N* is the total number of GR cells (i.e., *N* = *N*_*c*_ · *N*_GR_ = 51, 200). As a kernel function *K*_*h*_(*t*), we use a Gaussian kernel function of band width *h*:

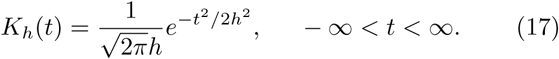

Throughout the paper, the band width *h* of *K*_*h*_(*t*) is 10 msec.

Figure 4(a) shows a raster plot of spikes of 10^3^ randomly chosen GR cells. At the beginning of trial stage (0 < *t* < 7 msec), all GR cells fire spikes due to the effect of strong transient CS signal of 200 Hz. In the remaining part of the trial stage (7 < *t* < 1000 msec), GR cells make random repetition of transitions between active and inactive states because of sustained CS signal of 30 Hz, and thus they seem to exhibit various spiking trains. Time passage from the CS onsets may be represented by the various firing patterns of GR cells, which will be explained in details in Figs. 5 and 6. In the break stage (1000 < *t* < 2000 msec), GR cells fire very sparse spikes. For simplicity, only the raster plot in the range of 1000 < *t* < 1100 msec is shown; the raster plot in a part of the preparatory stage (essentially the same as the break stage) (− 100 < *t* < 0 msec), just before the 1st trial stage, is also shown. Figure 4(b) shows the instantaneous whole-population spike rate *R*_GR_(*t*) in the whole population of GR cells. *R*_GR_(*t*) is basically in proportion to the transient and sustained CS inputs via MFs [see Figs. 1(b1)-1(b2)]. Thus, *R*_GR_(*t*) is sharply peaked in the beginning of the trial stage due to the strong transient CS, and then it becomes nearly flat in the remaining part of the trial stage where the sustained CS is present. However, due to the inhibitory effect of GO cells, the overall firing rates are uniformly lowered as follows. The time-averaged whole-population spike rates 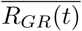 in the time intervals of 0 < *t* < 5 msec, 5 < *t* < 1000 msec, and 1000 < *t* < 2000 msec are 155.4 Hz, 32.5 Hz, and 3.4 Hz, respectively.

**FIG. 4:**
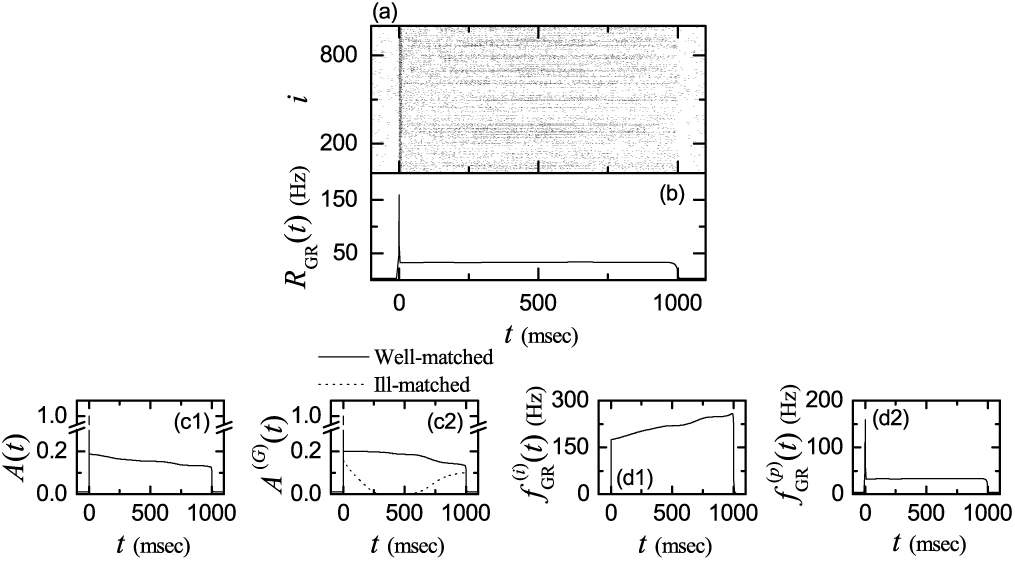
Firing activity of GR cells in an optimal case of *p*_*c*_ (connection probability from GO to GR cells) = 0.029. (a) Raster plots of spikes of 10^3^ randomly chosen GR cells and (b) instantaneous whole-population spike rate *R*_GR_(*t*) in the whole population of GR cells for the 1st step in the learning process for the EBC. Plots of the activation degrees (c1) *A*(*t*) in the whole population of GR cells and (c2) *A*^(*G*)^(*t*) in the *G* firing group [*G* (= *w*) : well-matched (solid curve) and *G* (= *i*) : ill-matched (dotted curve)]. Plots of (d1) instantaneous individual firing rate 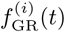 for the active GR cells and (d2) instantaneous population spike rate 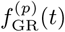 in the whole population of GR cells

**FIG. 5:**
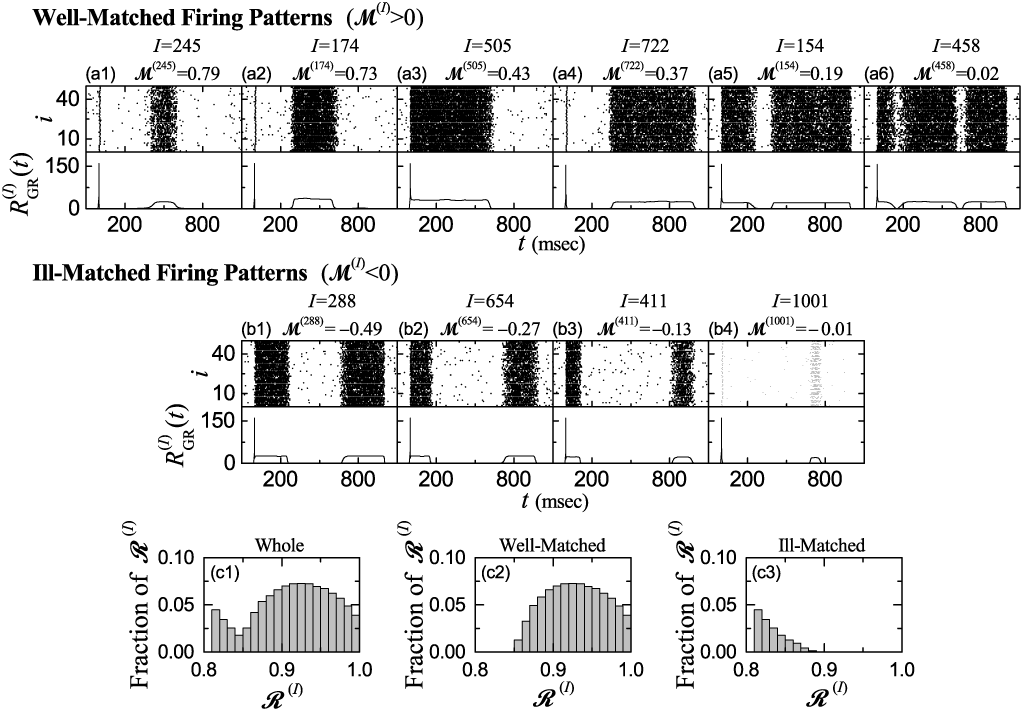
Various firing patterns in the GR clusters in an optimal case of 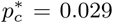. Raster plots of spikes and instantaneous cluster spike rates 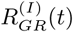 for various firing patterns. Six well-matched firing patterns in the *I*th GR clusters; *I* = (a1) 245, (a2) 174, (a3) 505, (a4) 722, (a5) 154, and (a6) 458. Four ill-matched firing patterns in the *I*th GR cluster; *I* = (b1) 288, (b2) 654, (b3) 411, and (b4) 1001. ℳ^(*I*)^ represents the matching index of the firing pattern in the *I*th GR cluster. Distribution of the reproducibility degree ℛ^(*I*)^ in the (c1) whole population and the (c2) well- and (c3) ill-matched firing groups. Bin size for the histograms in (c1)-(c3) is 0.01.

**FIG. 6:**
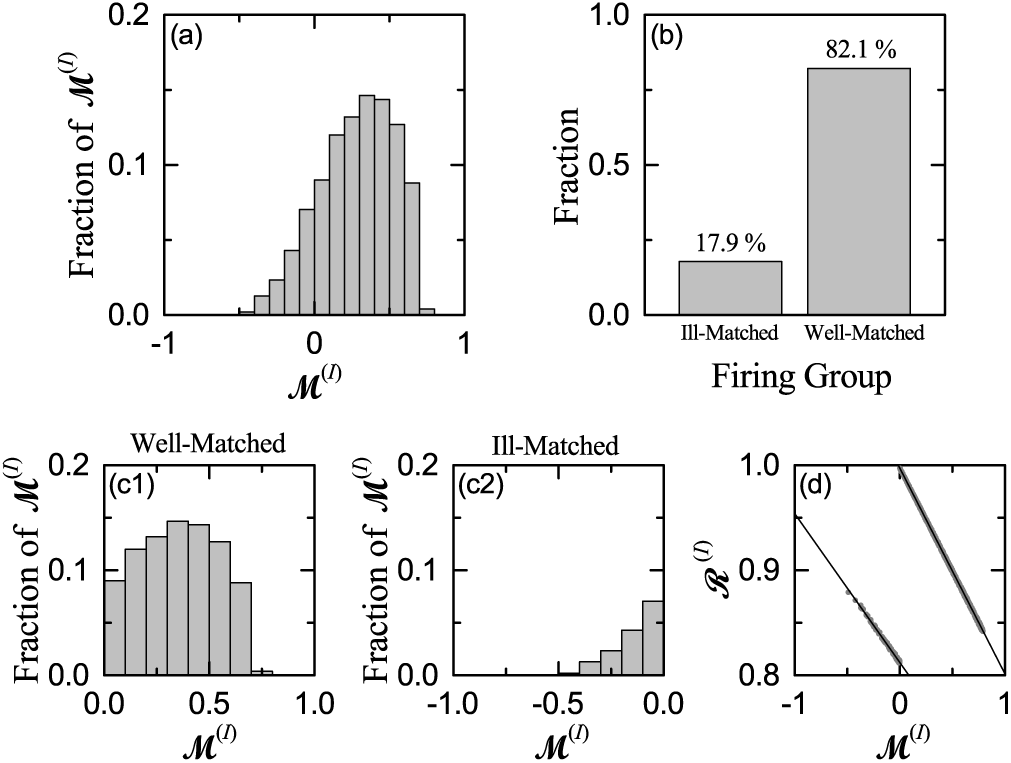
Characterization of various firing patterns in the GR clusters in an optimal case of 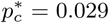. (a) Distribution of matching indices {ℳ^(*I*)^} in the whole population. (b) Fraction of well-matched and ill-matched firing groups. Distribution of matching indices {ℳ^(*I*)^} for the (c1) well- and (c2) ill-matched firing groups. Bin size for the histograms in (a) and in (c1)-(c3) is 0.1. (d) Plots of reproducibility degree ℛ^(*I*)^ versus ℳ^(*I*)^ in the well-matched (ℳ^(*I*)^ > 0) and the ill-matched (ℳ^(*I*)^ < 0) firing groups.

We next consider the activation degree of GR cells. To examine it, we divide the whole learning step into bins. In the beginning of the trial stage (0 < *t* < 10 msec), we divide the time interval into small bins (bin size: 1 msec) to properly take into consideration the effect of strong transient CS; the effect of transient CS seems to persist until the 7th bin. Then, in the remaining trial stage (10 < *t* < 1000 msec), to use wide bins (bin size: 10 msec) seems to be sufficient for considering the effect of sustained CS. Thus, we obtain the activation degree *A*_*i*_ for the active GR cells in the *i*th bin:

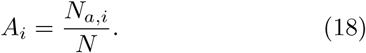

Here, *N*_*a,i*_ and *N* (= 51, 200) are the number of active GR cells in the *i*th bin and the total number of GR cells, respectively. Figure 4(c1) shows a plot of the activation degree *A*(*t*) in the whole population of GR cells. In the initial 7 bins (0 < *t* < 7 msec), *A* = 1 due to the effect of strong transient CS. In the presence of sustained In the break stage (1000 < *t* < 2000 msec), the time-averaged activation degree 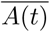 is 0.161. Hence, representation of time passage from the onsets of CS can be made in a sparse recoding scheme (i.e., MF inputs become more sparse via recoding in the granular layer). In the break stage (1000 < *t* < 2000 msec), the timeaveraged activation degree 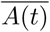 is 0.011 and small variations occur, which may be regarded as nearly “silent” stage, in comparison with the trial stage.

The whole population of GR cells may be decomposed into two types of well-matched and ill-matched firing groups; details will be given in Figs. 5 and 6. Firing patterns in the well-matched (ill-matched) firing group are well (ill) matched with the airpuff US signal. In this case, the activation degree 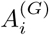 of active GR cells in the *i*th bin in the *G* firing group is given by:

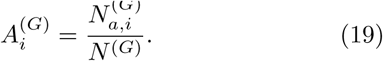

Here, 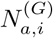 and *N*^(*G*)^ are the number of active GR cells in the *i*th bin and the total number of GR cells in the *G* firing group, respectively (*G* = *w* and *i* for the well-matched and the ill-matched firing groups, respectively). The number of clusters, belonging to the well- and the ill-matched firing groups are 841 and 183, respectively, and hence *N* ^(*w*)^ = 42, 050 and *N* ^(*i*)^ = 9, 150 because *N*_*GR*_ = 50 (number of GR cells in each cluster).

Figure 4(c2) shows plots of activation degree *A*^(*G*)^(*t*) in the well-matched (solid line) and the ill-matched (dotted curve) firing groups. In the beginning of the trial stage [i.e., in the initial 7 bins (0 < *t* < 7 msec)], *A*^(*G*)^ = 1, independently of the firing groups, due to the effect of strong transient CS. On the other hand, in the remaining trial stage (7 < *t* < 1000 msec) where the sustained CS is present, *A*^(*G*)^(*t*) varies, strongly depending on the type of firing groups. In the case of well-matched firing group, *A*^(*w*)^(*t*) decreases monotonically from 0.2 to 0.133, which is a little higher than *A*(*t*) in the whole population. In contrast, in the case of ill-matched firing group, *A*^(*i*)^(*t*) forms a well-shaped curve with a central “zero-bottom” with the time-averaged activation degree 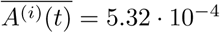 for 330 < *t* < 580 msec. Due to appearance of the central zero-bottom, contribution of the ill-matched firing group to *A*(*t*) in the whole population may be negligible in the range of 330 < *t* < 580 msec. In the break stage (1000 < *t* < 2000 msec), the time-averaged activation degree 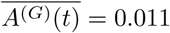 (*G* = *w* or *i*) with small variations, independently of the firing groups, which is the same as 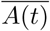 in the whole population.

In each *i*th bin, the contribution 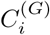 of each firing group to the activation degree *A*_*i*_ in the whole population is given by the product of the fraction *F* ^(*G*)^ and the activation degree 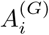 of the firing group:

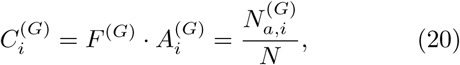

where *F* ^(*G*)^ = *N* ^(*G*)^*/N* ; *F* ^(*w*)^ = 0.821 (82.1%) and *F* ^(*i*)^ = 0.179 (17.9%) [see Fig. 6(b)]. Hence, we can easily get the contribution 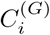 of each firing group by just multiplying 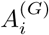 in Fig. 4(c2) with the fraction *F* ^(*G*)^. The sum of 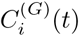 over the well- and the ill-matched firing groups is just the activation degree *A*_*i*_(*t*) in the whole population (i.e., 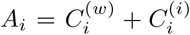). In this case, contribution 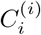 of the ill-matched firing group becomes small due to both low activation degree 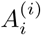 and small fraction *F* ^(*i*)^. Particularly, because of existence of the central zero-bottom, contribution 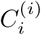 is negligibly small in the middle (330 < *t* < 580 msec) of the trial stage.

In the whole population, the activation degree *A*(*t*) showing decreasing tendency is in contrast to the instantaneous whole-population spike rate *R*_GR_(*t*) which is flat in the trial stage. To understand this discrepancy, we consider the bin-averaged instantaneous individual firing rate 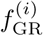 of active GR cells:

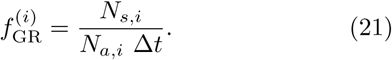

Here, *N*_*s,i*_ is the number of spikes of GR cells in the *i*th bin, *N*_*a,i*_ is the number of active GR cells in the *i*th bin, and Δ*t* is the bin size. Figure 4(d1) shows a plot of 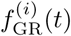 for the active GR cells. In the initial 7 bins (0 < *t* < 7 msec) of the trial stage where *A*(*t*) = 1, 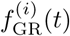 decreases very slowly from 155.6 to 155.3 Hz with the time *t* (i.e., the values of 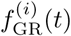 are nearly the same). In the remaining part (7 < *t* < 1000 msec) of the trial stage, 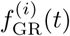 increases monotonically from 173 to 258 Hz. In this case, the bin-averaged instantaneous population spike rate 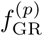 is given by the product of the activation degree *A*_*i*_ of Eq. (18) and the instantaneous individual firing rate 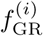 of Eq. (21):

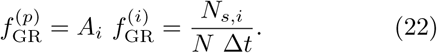

Figure 4(d2) shows a plot of the instantaneous population spike rate 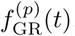. It is flat except for the sharp peak in the beginning of the trial stage, as in the case of *R*_GR_(*t*). We note that both 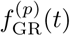 and *R*_GR_(*t*) correspond to bin-based estimate and kernel-based smooth estimate for the instantaneous whole-population spike rate of the GR cells, respectively [76]. Although the activation degree *A*(*t*) of GR cells decreases with *t*, their population spike rate keeps the flatness (i.e., 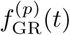 becomes flat), because of the increase in the individual firing rate 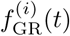. As a result, the bin-averaged instantaneous population spike rate 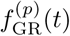 in Fig. 4(d2) becomes essentially equal to the instantaneous whole-population spike rate *R*_GR_(*t*) in Fig. 4(b).

### B. Dynamical Classification and Dynamical Origin of Various Firing Patterns in The GR Clusters

There exist *N*_*C*_ (= 2^10^) GR clusters in the whole population. *N*_*GR*_ (= 50) GR cells in each GR cluster share the same inhibitory and excitatory inputs via their dendrites which synaptically contact the four glomeruli (i.e., terminals of MFs) at both ends of the GR cluster [see Fig. 2(b)]. Nearby inhibitory GO cell axons innervate the four glomeruli. Due to the shared inputs, GR cells in each GR cluster exhibit similar firing behaviors.

As in the case of *R*_GR_(*t*) in Eq. (16), the firing activity of the *I*th GR cluster is described in terms of its instantaneous cluster spike rate 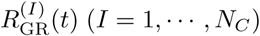

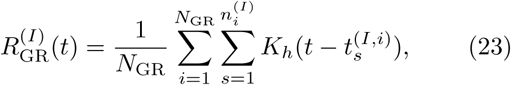

where 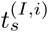 is the *s*th firing time of the *i*th GR cell in the *I*th GR cluster and 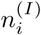 is the total number of spikes for the *i*th GR cell in the *I*th GR cluster.

We introduce the matching index ℳ^(*I*)^ of each GR cluster between the firing behavior 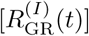 of each *I*th GR cluster and the airpuff US signal *f*_US_(*t*) for the desired timing [see Fig.1(c)]. The matching index ℳ^(*I*)^ is given by the cross-correlation at the zero-time lag [i.e., 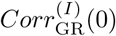] between 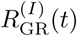 and *f*_US_(*t*):

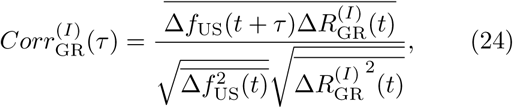

where 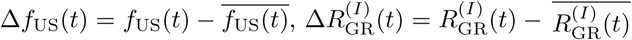, and the overline denotes the time average. We note that ℳ^(*I*)^ represents well the phase difference between the firing patterns 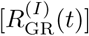 of GR clusters and the US signal [*f*_US_(*t*)].

Figure 5 shows various firing patterns of GR clusters. This type of variety results from inhibitory coordination of GO cells on the firing activity of GR cells in the GRGO feedback loop in the granular layer. Time passage from the CS onsets may be well represented by the various firing patterns of GR clusters because MF inputs become less similar (i.e., more orthogonal) to each other through recoding in the granular layer.

Six examples for the well-matched firing patterns in the *I*th (*I* = 245, 174, 505, 722, 154, and 458) GR clusters are given in Figs. 5(a1)-5(a6), respectively. Raster plot of spikes of *N*_GR_ (= 50) GR cells and the corresponding instantaneous cluster spike rate 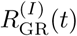 are shown, along with the value of the matching index ℳ^(*I*)^ in each case of the *I*th GR cluster. In all these cases, the instantaneous cluster spike rates 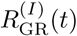 are well-matched with the US signal *f*_US_(*t*), and hence these well-matched GR clusters have positive matching indices (i.e., ℳ^(*I*)^ > 0).

In the 1st case of *I* = 245 with the highest ℳ^(*I*)^ (= 0.79), 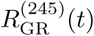 is strongly localized around the middle of the trial stage (i.e. a central band of spikes is formed around *t* = 500 msec), and hence it is the most well-matched with the US signal *f*_US_(*t*). In the 2nd case of *I* = 174 with ℳ^(*I*)^ = 0.73, 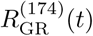 is also localized around *t* = 500 msec, but its central firing band spreads a little more to the left side, in comparison with the case of *I* = 245. Hence, its matching index relative to *f*_US_(*t*) is a little decreased.

We note that LTD at the PF-PC synapses occurs within an effective range of 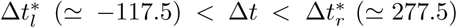 (see Fig. 3). Here, Δ*t* [= *t*_CF_ (CF activation time) - *t*_PF_ (PF activation time)] is the relative time difference between the firing times of the error-teaching instructor CF and the variously-recoded student PF. The CF activation occurs approximately at *t*_CF_ = 500 msec due to the strong brief US (strongly localized at *t* = 500 msec). Then, LTD may occur when the PF activation time *t*_PF_ lies in an effective LTD range of 222.5 msec < *t*_PF_ < 617.5 msec. In the above two cases of GR clusters (*I* = 245 and 174) with higher ℳ^(*I*)^, their PF signals (corresponding to axons of the GR cells) are strongly depressed by the US instructor signal because most parts of their firing bands are well localized in the effective LTD range.

We next consider the 3rd and the 4th cases of the *I*th GR cluster (*I* = 505 and 722) with intermediate ℳ^(*I*)^. In the cases of *I* = 505 (722), the firing band in the raster plot extends to the left (right) until *t* ≃ 0 (1000) msec. Thus, big left- and right-extended firing bands appear for *I* = 505 and 722, respectively. Some part of this big firing band lies inside the effective LTD range where LTD occurs in conjunction with the CF firing. On the other hand, its remaining part lies outside the effective LTD range, and hence LTP occurs for the PF firings alone without association with the CF signal.

We also consider the case of lower ℳ^(*I*)^ for *I* = 154 and 458 (i.e., the 5th and the 6th cases). In both cases, they have tendency to fill the raster plots with more spikes via appearance of two or more firing bands. Thus, some central part of these bands lies inside the effective LTD range where LTD occurs. In contrast, LTP occurs in the other left and right parts of the firing bands because they lie outside the effective LTD range; in comparison with the case of intermediate ℳ^(*I*)^, LTP region is extended. In this way, as ℳ^(*I*)^ is decreased toward the zero, the raster plot tends to be filled with more spikes (constituting firing bands), and hence the region where LTP occurs is extended.

In addition to the well-matched firing patterns, ill-matched firing patterns with negative matching indices (i.e., ℳ^(*I*)^ < 0) also appear. Four examples for the ill-matched firing patterns in the *I*th (*I* = 288, 654, 411, and 1001) GR clusters are given in Figs. 5(b1)-5(b4), respectively. We first consider the case of *I* = 288 with the lowest ℳ^(*I*)^ (= −0.49) (i.e., its magnitude |ℳ^(*I*)^|: largest). This lowest case corresponds to the “opposite” case of the highest one for *I* = 245 with ℳ^(*I*)^ = 0.79 in the following sense. A central gap with negligibly small number of spikes (i.e., practically no spikes) appears around *t* = 500 msec, in contrast to the highest case where a central firing band exists. Hence, in this lowest case, occurrence of LTD in the central gap may be practically negligible. On the other hand, mainly LTP occurs in the left and right firing bands, most of which lie outside the effective LTD range. The right firing band lies completely outside the effective LTD range, and hence no LTD occurs. The width of the central gap is larger than the width of the effective LTD range. However, since the gap is shifted a little to the right, a small part near the right boundary (*t* ≃ 261 msec) of the left firing band overlaps with a small region near the left boundary (*t* ≃ 222.5 msec) of the effective LTD region. In this small overlapped region of 239 < Δ*t* < 277.5 msec, the values of the synaptic modification Δ*J*_LTD_ (i.e., the average synaptic modification ⟨Δ*J*_LTD_ ⟩ ≃0.031) are very small, and hence very weak LTD may occur.

As the magnitude |ℳ^(*I*)^| is decreased, the central gaps becomes widened, and the widths of the left and the right firing bands also get decreased, as shown in the cases of *I* = 654, 411, and 1001. In these cases, the two left and right firing bands lie completely outside the effective LTD range, and hence only LTP occurs for the PF signals alone without conjunction with the CF signal. In this way, as |ℳ^(*I*)^| approaches the zero from the negative side, spikes become more and more sparse, which is in contrast to the well-matched case where more and more spikes fill the raster plot as ℳ^(*I*)^ goes to the zero from the positive side.

The above firing patterns are shown in the 1st learning step [consisting of the 1st trial stage (0 < *t* < 1000 msec) and the 1st break stage (1000 < *t* < 2000 msec)]. For simplicity, they are shown in the range of 1000 < *t* < 1100 msec in the break stage, and a part of the preliminary stage (− 100 < *t* < 0 msec), preceding the 1st learning step, is also shown. We examine the reproducibility of the firing patterns across the learning steps. To this end, we consider the cross-correlation between the instantaneous cluster spike rates 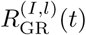 in the *I*th GR cluster for the *k*th (*l* = *k*) and the (*k*+1)th (*l* = *k*+1) learning steps;

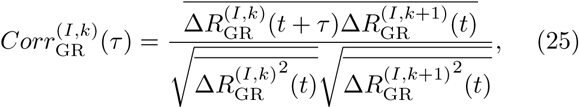

where 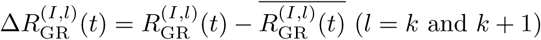 and the overline represents the time average. Then, the reproducibility degree *R*^(*I*)^ of the *I*th GR cluster is given by the average value of the cross-correlations at the zero-time lag between the instantaneous cluster spike rates 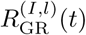 for the successive learning steps:

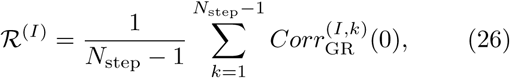

where *N*_step_ is the total number of learning steps. Here, we consider the case of *N*_step_ = 100.

Figure 5(c1) shows the distribution of the reproducibility degrees ℛ^(*I*)^ for the whole GR clusters. Double peaks appear; large broad peak and small sharp peaks at ℛ^(*I*)^ = 0.925 and 0.815, respectively. The range of ℛ^(*I*)^ is (0.812, 0.997). Hence, the firing patterns are highly reproducible across the learning steps. Figures 5(c2) and 5(c3) also show the distributions of ℛ^(*I*)^ of the GR clusters in the well- and the ill-matched firing groups, respectively. In the case of well-matched firing group, the distribution of ℛ^(*I*)^ has a broad peak and its range is (0.842, 0.997). On the other hand, in the ill-matched case, the distribution decreases from its maximum at ℛ^(*I*)^ = 0.815, and its range is (0.812, 0.879). We note that the average values of {ℛ^(*I*)^} in the well- and the ill-matched firing group are 0.927 and 0.828, respectively. Hence, on average, the firing patterns of the GR clusters in the well-matched firing group may be more reproducible than those in the ill-matched firing group, because the average individual firing rate in the well-matched firing group is higher than that in the ill-matched firing group.

Results on characterization of the various well- and ill-matched firing patterns are given in Fig. 6. Figure 6(a) shows the plot of the fraction of matching indices ℳ^(*I*)^ in the whole GR clusters. ℳ^(*I*)^ increases slowly from the negative value to the peak at 0.35, and then it decreases rapidly. For this distribution of {ℳ^(*I*)^}, the range is (−0.49, 0.79), the mean is 0.3331, and the standard deviation is 0.6135. Then, we obtain the variety 𝒱 degree for the firing patterns 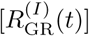 of all the GR clusters:

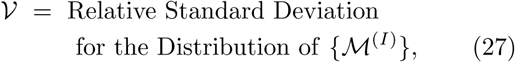

where the relative standard deviation is just the standard deviation divided by the mean. In the optimal case of 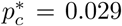, 𝒱 ^*^ ≃ 1.842, which is just a quantitative measure for the various recoding made through feedback cooperation between the GR and the GO cells in the granular layer. It will be seen that 𝒱 ^*^ is just the maximum in Fig. 16(b) for the plot of 𝒱 versus *p*_*c*_. Hence, firing patterns of the GR clusters at 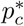 is the most various.

We decompose the whole GR clusters into the well-matched ({ℳ^(*I*)^} > 0) and the ill-matched ({ℳ^(*I*)^} < 0) firing groups. Figure 6(b) shows the fraction of firing groups. The well-matched firing group is a major one with fraction 82.1%, while the ill-matched firing group is a minor one with fraction 17.9%. In this case, the firing-group ratio ℛ_sp_, given by the ratio of the fraction of the well-matched firing group to that of the ill-matched firing group is 4.59. For this firing-group ratio, firing patterns of the GR clusters are the most various.

Figures 6(c1)-6(c2) also show the plots of matching indices ℳ^(*I*)^ of the GR clusters in the well- and the ill-matched firing groups, respectively. In the case of well-matched firing group, the distribution of ℳ^(*I*)^ with a peak at 0.35 has only positive values in the range of (0.0, 0.79), and its mean and standard deviations are 0.428 and 0.372, respectively. On the other hand, in the case of the ill-matched firing group, the distribution of ℳ^(*I*)^ with a maximum at -0.05 has only negative values in the range of (−0.49, 0.0), and its mean and standard deviations are -0.104 and 0.129, respectively. In this case, ℳ^(*I*)^ increases slowly to the maximum. As will be seen in the next subsection, these well- and the ill-matched firing groups play their own roles in the synaptic plasticity at PF-PC synapses and the subsequent learning process for the EBC, respectively.

We also examine the correlation between the atching index ℳ^(*I*)^ and the reproducibility degree ℛ^(*I*)^. Figure 6(d) show the plots of ℳ^(*I*)^ versus ℛ^(*I*)^ in the well- and the ill-matched firing groups. In both cases, there appear strong negative correlations between ℳ^(*I*)^ and ℛ^(*I*)^; for the well-matched (ill-matched) firing group, the Pearson’s correlation coefficient is *r* = − 0.9999 (− 0.9978). When left-right reflections are made on Figs. 6(c1)-6(c2), shapes of the reflected ones are similar to the shapes of Figs. 5(c2)-5(c3), respectively, which implies the negative correlation between ℳ^(*I*)^ and ℛ^(*I*)^ in each firing group.As shown in Figs. 5(a1)-5(a6), as ℳ^(*I*)^ decreases to the zero from the positive side, the raster plot tends to be filled with more spikes due to increased individual firing rates, which leads to increase in ℛ^(*I*)^. On the other hand, as ℳ^(*I*)^ increases to the zero from the negative side, the raster plot of spikes tends to be more sparse because of decreased individual firing rates [see Figs. 5(b1)-5(b4)], which results in decrease in ℛ^(*I*)^. Consequently, there appears a gap at the limit ℳ^(*I*)^ = 0.

Finally, we study the dynamical origin of various firing patterns in the *I*th GR clusters. As examples, we consider two well-matched firing patterns for *I* = 245 and 722 [see the firing patterns in Figs. 5(a1) and 5(a4)] and two ill-matched firing patterns for *I* = 288 and 654 [see the firing patterns Figs. 5(b1) and 5(b2)]. In Fig. 7, (a1)-(a4) correspond to the cases of *I* =245, 722, 288, and 654, respectively.

**FIG. 7:**
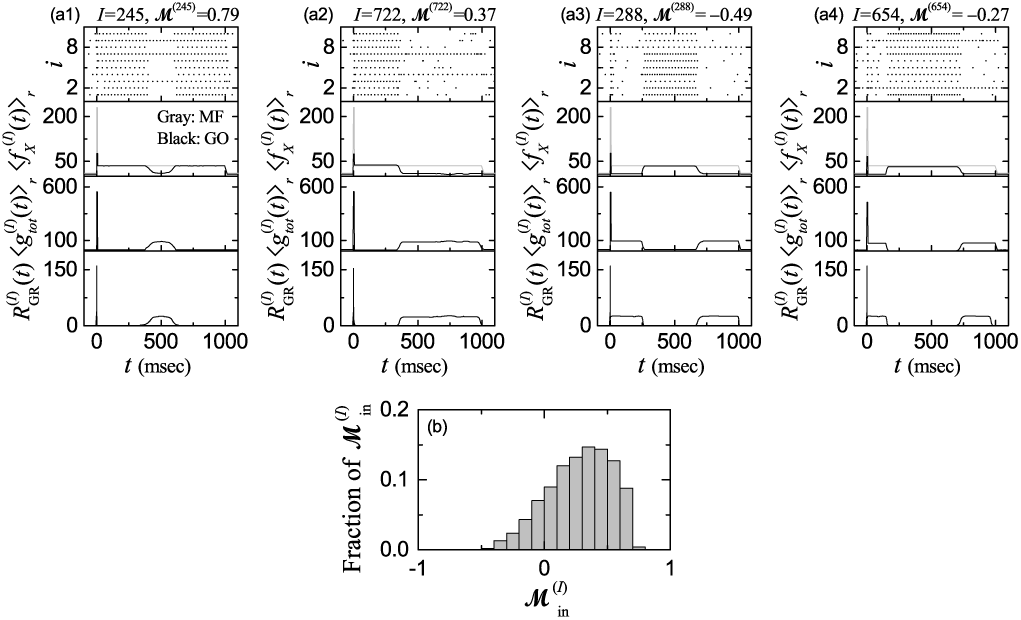
Dynamical origin of various firing patterns in the GR clusters in an optimal case of 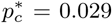. Well-matched firing patterns for *I*= (a1) 245 and (a2) 722 and ill-matched firing patterns for *I*= (a3) 288 and (a4) 654. In (a1)-(a4), top panel: raster plots of spikes in the sub-population of pre-synaptic GO cells innervating the *I*th GR cluster, 2nd panel: plots of 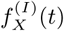 : bin-averaged instantaneous spike rates of the MF signals (*X* = MF) into the *I*th GR cluster (gray line) and bin-averaged instantaneous sub-population of pre-synaptic GO cells (*X* = GO) innervating the *I*th GR cluster (black line); ⟨*…* ⟩_*r*_ represents the realization average (number of realizations is 100), 3rd panel: time course of 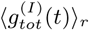: conductance of total synaptic inputs (including both the excitatory and inhibitory inputs) into the *I*th GR cluster, and bottom panel: plots of 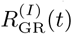 : instantaneous cluster spike rate in the *I*th GR cluster. (b) Distribution of matching indices 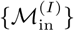 for the conductances of total synaptic inputs into the GR clusters.

Various recodings for the MF signals are made in the GR layer, consisting of excitatory GR and inhibitory GO cells (i.e., in the GR-GO cell feedback loop). Thus, firing activities of GR cells are determined by two types of synaptic input currents (i.e., excitatory synaptic inputs via MF signals and inhibitory synaptic inputs from randomly connected GO cells). Then, investigations on the dynamical origin of various firing patterns of the GR clusters are made via analysis of total synaptic inputs into the GR clusters. As in Eq. (6), synaptic current is given by the product of synaptic conductance *g* and potential difference. In this case, synaptic conductance determines the time-course of synaptic current. Hence, it is sufficient to consider the time-course of synaptic conductance. The synaptic conductance *g* is given by the product of synaptic strength per synapse, the number of synapses *M*_syn_, and the fraction *s* of open (post-synaptic) ion channels [see Eq. (7)]. Here, the synaptic strength per synapse is given by the product of maximum synaptic conductance 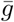 and synaptic weight *J*, and the time-course of *s*(*t*) is given by a summation for exponential-decay functions over pre-synaptic spikes, as shown in Eqs. (7) and (8).

Here, we make an approximation of the fraction *s*(*t*) of open ion channels (i.e., contributions of summed effects of pre-synaptic spikes) by the bin-averaged spike rate 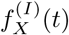 of pre-synaptic neurons (*X* = MF and GO); 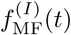 is the bin-averaged spike rate of the MF signals into the *I*th GR cluster and 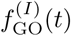 is the bin-averaged spike rate of the pre-synaptic GO cells innervating the *I*th GR cluster. In the case of MF signal, we get:

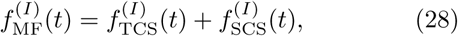

where 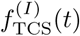 and 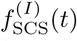 are the bin-averaged spike rates of the transient and the sustained CS signals, respectively.

Then, the conductance 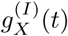 of synaptic input from *X* (=MF or GO) into the *I*th GR cluster (*I* = 1, *…, N*_*C*_) is given by:

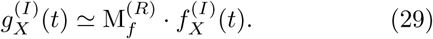

Here, the multiplication factor 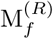 [= maximum synaptic conductance 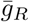 × synaptic weight *J* ^(GR,*X*)^ × number of synapses 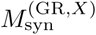] varies depending on *X* and the receptor *R* on the post-synaptic GR cells. In the case of excitatory synaptic currents into the *I*th GR cluster with AMPA receptors via TCS or SCS MF signal, 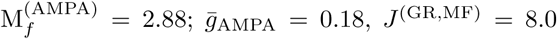, and 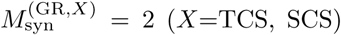. On the other hand, in the case of the *I*th GR cluster with NMDA receptors, 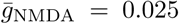, and hence 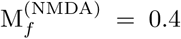, which is much less than 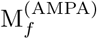. For the inhibitory synaptic current from pre-synaptic GO cells to the *I*th GR cluster with GABA receptors, 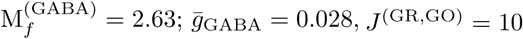, and 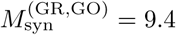. Then, the conductance 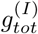 of total synaptic inputs (including both the excitatory and the inhibitory inputs) into the *I*th GR cluster is given by:

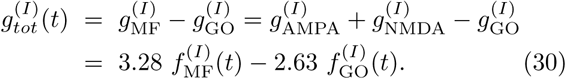

In Figs. 7(a1)-7(a4), the top panels show the raster plots of spikes in the sub-populations of pre-synaptic GO cells innervating the *I*th GR clusters. From these raster plots, bin-averaged (sub-population) spike rates 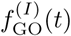 may be obtained. The bin-averaged spike rate of pre-synaptic GO cells in the *i*th bin is given by 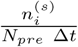, where 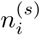 is the number of spikes in the *i*th bin, Δ*t* is the bin size, and *N*_*pre*_ (=10) is the number of pre-synaptic GO cells. As in Fig. 4, in the beginning of the trial stage (0 < *t* < 10 msec), we employ the small bin-size of Δ*t* = 1 msec to properly take into consideration the effect of strong transient CS, and in the remaining trial stage (10 < *t* < 1000 msec), we use the wide bin-size of Δ*t* = 10 msec for considering the effect of sustained CS. Through an average over 100 realizations, we obtain the realization-averaged (bin-averaged) spike rate of presynaptic GO cells 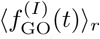 because *N*_*pre*_ (= 10) is small; ⟨…⟩_*r*_ represent a realization-average. The 2nd panels show 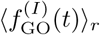 (black line) and 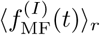 (gray line). We note 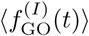 varies depending on I, while 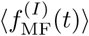 is independent of *I*. Then, we obtain the realization-averaged conductance 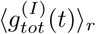 of total synaptic inputs in Eq. (30), which is shown in the 3rd panels.

We note that the shapes of 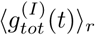 (corresponding to the total input into the *I*th GR cluster) in the 3rd panels are nearly the same as those of 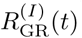 (corresponding to the output of the *I*th GR cluster) in the bottom panels. It is thus expected that well-matched (ill-matched) inputs into the GR clusters may lead to generation of well-matched (ill-matched) outputs (i.e., responses) in the GR clusters. To confirm this point clearly, as in case of the firing patterns 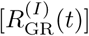 in the GR clusters, we introduce the matching index for the total synaptic input of the *I*th GR cluster between 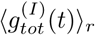 (conductance of total synaptic input into the *I*th GR cluster) and the (airpuff) US signal *f*_US_(*t*) for the desired timing. Similar to the matching index ℳ^(*I*)^ for the firing patterns (i.e. outputs) in the *I*th GR cluster [see Eq. (24)], the matching index 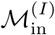 for the total synaptic input is given by the crosscorrelation at the zero-time lag (i.e., 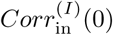) between 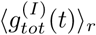 and *f*_US_(*t*):

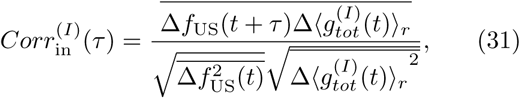

where 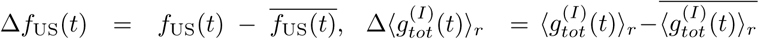, and the overline represents the time average. Thus, we have two types of matching indices, *ℓ*^(*I*)^ [output matching index: given by 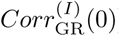] and 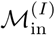 [input matching index: given by 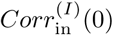] for the output and the input in the *I*th GR cluster, respectively.

Figure 7(b) shows the plot of fraction of input matching indices 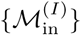 in the whole GR clusters. We note that the distribution of input matching indices in Fig. 7(b) is nearly the same as that of output matching indices in Fig. 6(a). 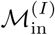 increases slowly from the negative value to the peak at 0.35, and then it decreases rapidly. In this distribution of 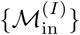, the range is (−0.49, 0.79), the mean is 0.3332, and the standard deviation is 0.6137. Then, we get the variety degree 𝒱_in_ for the total synaptic inputs 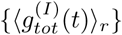 of all the GR clusters:

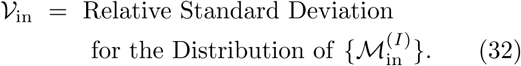

Hence, 𝒱_in_ ≃ 1.842 for the synaptic inputs, which is nearly the same as 𝒱^*^ (≃ 1.842) for the firing patterns of GR cells. Consequently, various synaptic inputs into the GR clusters results in generation of various outputs (i.e., firing patterns) of the GR cells.

### C. Influence of Various Temporal Recoding on Synaptic Plasticity at PF-PC Synapses

Based on dynamical classification of firing patterns of GR clusters, we study the influence of various temporal recoding in the GR clusters on synaptic plasticity at PFPC synapses. As shown in the preceding subsection, MF context input signals for the EBC are variously recoded in the granular layer (corresponding to the input layer of the cerebellar cortex). The variously-recoded well- and ill-matched PF (student) signals (coming from the GR cells) are fed into the PCs (i.e., principal cells of the cerebellar cortex) and the BCs in the Purkinje-molecular layer. The PCs also receive well-matched error-teaching CF (instructor) signals from the IO, together with the inhibitory inputs from the BCs. Then, the synaptic weights at the PF-PC synapses vary depending on the matching degree between the PF and the CF signals.

We first consider the change in normalized synaptic weights 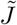 of active PF-PC synapses during the learning trials in the optimal case of 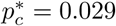;

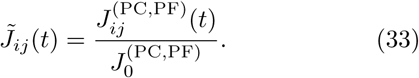

Here, the initial synaptic strength 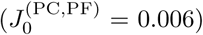 is the same for all PF-PC synapses. Figures 8(a1)-8(a9) show trial-evolution of distribution of 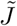 of active PF-PC synapses. As the learning trial is increased, normalized synaptic weights 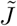 change due to synaptic plasticity at PF-PC synapses. We note that the distribution of 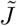 in each trial is composed of two markedly separated structures (i.e., a combination of separate top horizontal line with a central gap and lower band). Here, the top horizontal line with a central gap has no essential change with the trials, while the lower bands come down with the trials and their vertical widths increase. This kind of distribution of 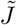 becomes saturated at about the 250th trial.

**FIG. 8:**
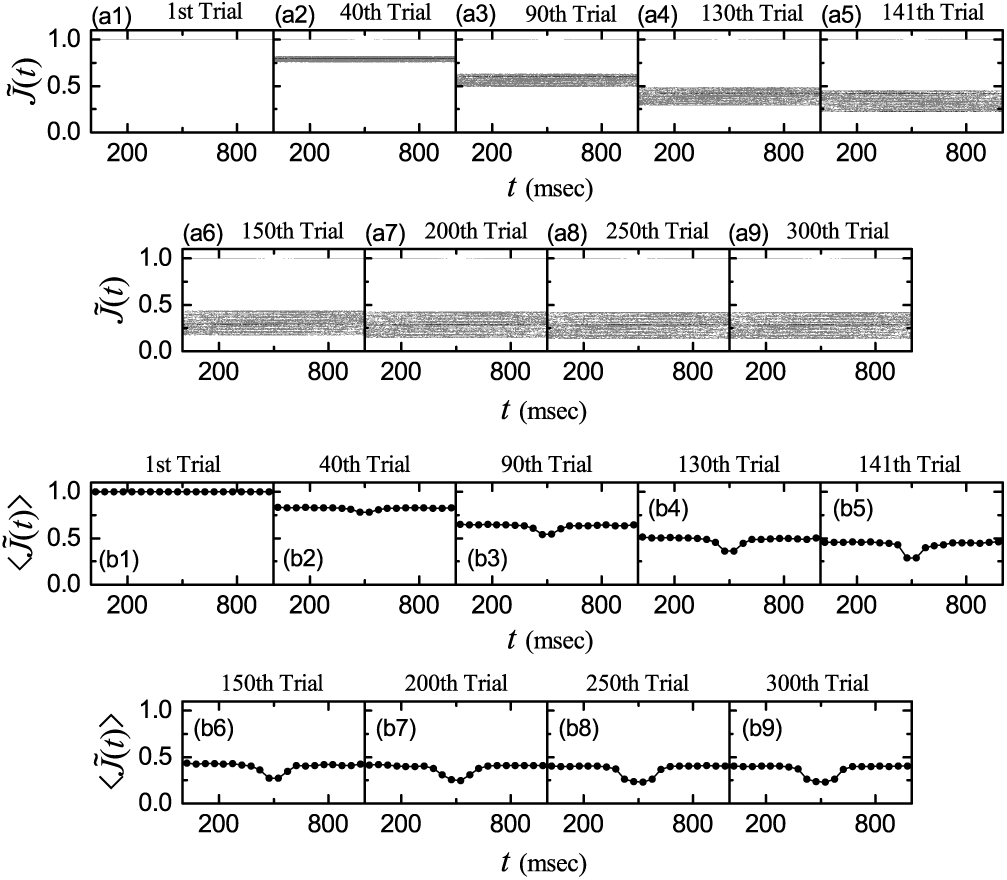
Change in synaptic weights of active PF-PC synapses during learning trials in the optimal case of 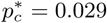. (a1)-(a9) Trial-evolution of distribution of normalized synaptic weights 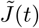 of active PF signals. (b1)-(b9) Trial-evolution of bin-averaged (normalized) synaptic weights 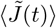 of active PF signals. Bin size: Δ*t* = 50 msec.

The top horizontal line with a central gap arises from the ill-matched firing group (with negative matching indices). In the case of ill-matched PF signals, practically no LTD occurs because most of them have no associations with the error-teaching CF signals which are strongly localized in the middle of trial (i.e., near *t* = 500 msec). As shown in Fig. 4(c2), the activation degree *A*^(*i*)^(*t*) (denoted by the dotted line) of GR cells in the ill-matched firing group has a central “zero-bottom” where *A*^(*i*)^(*t*) ≃ 0 (i.e., negligibly small number of spikes in the middle part of trial). In the initial and the final parts of the trial (outside the middle part), practically no LTD takes place due to no practical conjunctions with the strongly-localized CF signals. Thus, the normalized synaptic weights 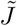 of the active GR cells in the ill-matched firing group forms the top horizontal line with a central gap which is nearly invariant with the trial.

On the other hand, lower bands arise from the well-matched firing group (with positive matching indices). In the case of well-matched PF signals, they are strongly depressed by the error-teaching CF signals (i.e., strong LTD occurs) in each trial due to good association between the well-matched PF and CF signals. As a result, a lower band is formed, it comes down with the trial, and eventually becomes saturated.

To more clearly examine the above trial evolutions, we obtain the bin-averaged (normalized) synaptic weight in each *i*th bin (bin size: Δ*t* = 50 msec):

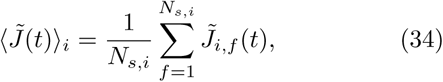

where 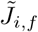 is the normalized synaptic weight of the *f* th active PF signal in the *i*th bin, and *N*_*s,i*_ is the total number of active PF signals in the *i*th bin. Figures 8(b1)-8(b9) show trial-evolution of bin-averaged (normalized) synaptic weights 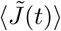 of active PF signals. In each trial, 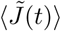 forms a step-well-shaped curve. As the trial is increased, the step-well curve comes down, its width and depth increase, and saturation seems to occur at about the 250th cycle.

We also obtain the trial-averaged mean 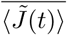 via time average of 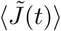 over a trial:

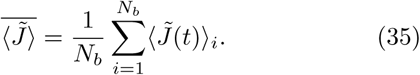

Here, *N*_*b*_ is the number of bins for trial averaging, and the overbar represents the time average. Figures 9(a) and 9(b) show plots of the trial-averaged mean 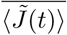 and the modulation [=(maximum - minimum)/2] ℳ_*J*_ for 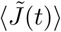 versus trial. The trial-averaged mean 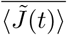 decreases monotonically from 1.0 due to LTD at PF-PC synapses, and it becomes saturated at 0.367 at about the 250th trial.

**FIG. 9:**
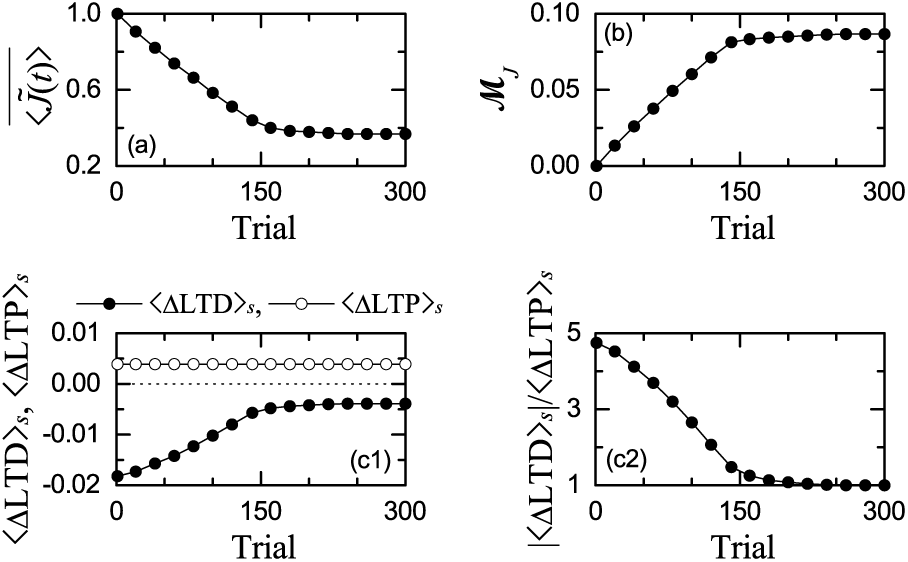
Plots of (a) trial-averaged mean 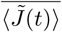 and (b) modulation ℳ_*J*_ for the bin-averaged (normalized) synaptic weights 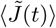 in Figs. 8(b1)-8(b9) versus trial. Dynamical balance between ΔLTD and ΔLTP during learning trials. Plots of (c1) trial-summed ΔLTD [⟨ΔLTD⟩_*s*_ (solid circles)] and ΔLTP [(ΔLTP)_*s*_ (open circles)] of active PF signals and (c2) ratio of |⟨ΔLTD⟩_*s*_| to (ΔLTP)_*s*_ versus trial.

However, strength of the LTD varies depending on the parts of the trial. In the middle part without practical contribution of ill-matched firing group, strong LTD occurs, due to contribution of only well-matched active PF signals. On the other hand, at the initial and the final parts, somewhat less LTD takes place, because both the ill-matched firing group (with practically no LTD) and the well-matched firing group make contributions together. Consequently, with increasing trial, the middle-stage part comes down more rapidly than the initial and final parts. Hence, the modulation ℓ_*J*_ increases monotonically from 0, and it gets saturated at 0.0867 at about the 250th trial.

We investigate the dynamical origin for the saturation of trial-averaged mean 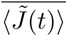 in Fig. 9(a). As explained in the Subsec. II E, a major LTD occurs when a CF firing makes association with earlier PF signals [see the 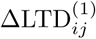 in Eq. (12)], while a minor LTD takes place when the CF firing makes conjunction with later CF signals [see the 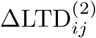 in Eq. (13)]. We consider the trial-summed ΔLTD (⟨ΔLTD ⟩_*s*_) in each trial, given by the sum of ΔLTD_*ij*_ occurring at all active PF-PC synapses during the whole trial (0 < *t* < 1000 msec), where 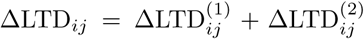. Similarly, we also consider the trial-summed ΔLTP (⟨ΔLTP ⟩_*s*_) in each trial, given by the sum of ΔLTP_*ij*_ occurring at all active PF-PC synapses during the whole trial. Here, LTP occurs for the PF signals alone without association with CF signals; ΔLTP_*ij*_ is given in Eq. (14).

Figures 9(c1) shows trial-evolution of trial-summed ΔLTD [⟨ΔLTD ⟩_*s*_ (solid circles)] and ΔLTP [⟨ΔLTP ⟩_*s*_ (open circles)] occurring at all active PF-PC synapses. Here, ⟨ΔLTP⟩ _*s*_ (≃0.00386), independently of the trial. In the beginning trials, ΔLTD _*s*_ (i.e., magnitude of ⟨ΔLTD ⟩_*s*_) is larger than ΔLTP _*s*_, and hence ΔLTD is dominant. However, with increasing the trial, | ⟨ΔLTD ⟩_*s*_ | decreases monotonically, and becomes saturated at 0.00386 at about the 250th trial. Thus, the saturated value of | ⟨ΔLTD ⟩_*s*_ | becomes the same as that of ⟨ΔLTP ⟩_*s*_. This process may be well seen in Fig. 9(c2). With increasing the trial, the ratio of | ⟨ΔLTD ⟩_*s*_ | _*s*_ to ⟨ΔLTP⟩ _*s*_ decreases from 4.71, and it converges to 1 at about the 250th trial. Thus, trial-level balance between ΔLTD and ΔLTP occurs [i.e., | ⟨ΔLTD ⟩_*s*_ | = ⟨ΔLTP ⟩_*s*_ (0.00386)] at about the 250th trial, which results in the saturation of trial-averaged mean 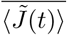.

### D. Influence of PF-PC Synaptic Plasticity on Subsequent Learning Process in The PC-CN-IO System

As a result of various recoding in the GR clusters, well- and ill-matched firing groups appear. In the case of well-matched PF signals, they are strongly depressed by the CF signals due to good association between the PF and CF signals. On the other hand, in the case of ill-matched PF signals, practically no LTD occurs because most of them have no conjunctions with the error-teaching CF signals. In this subsection, we investigate the influence of this kind of effective PF-PC synaptic plasticity on the subsequent learning process in the PC-CN-IO system.

Figure 10 shows change in firing activity of PCs during learning trial in the optimal case of 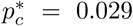. Trial-evolutions of raster plots of spikes of 16 PCs and the corresponding instantaneous population spike rates *R*_PC_(*t*) are shown in Figs. 10(a1)-10(a9) and Figs. 10(b1)-10(b9), respectively. Realization-averaged smooth instantaneous population spike rates ⟨*R*_PC_(*t*) _*r*_⟩ are also shown in Figs. 10(c1)-10(c9). Here, ⟨*…* _*r*_ ⟩ denotes realization average and the number of realizations is 100. ⟨*R*_PC_(*t*)⟨_*r*_ seems to be saturated at about the 250th cycle.

**FIG. 10:**
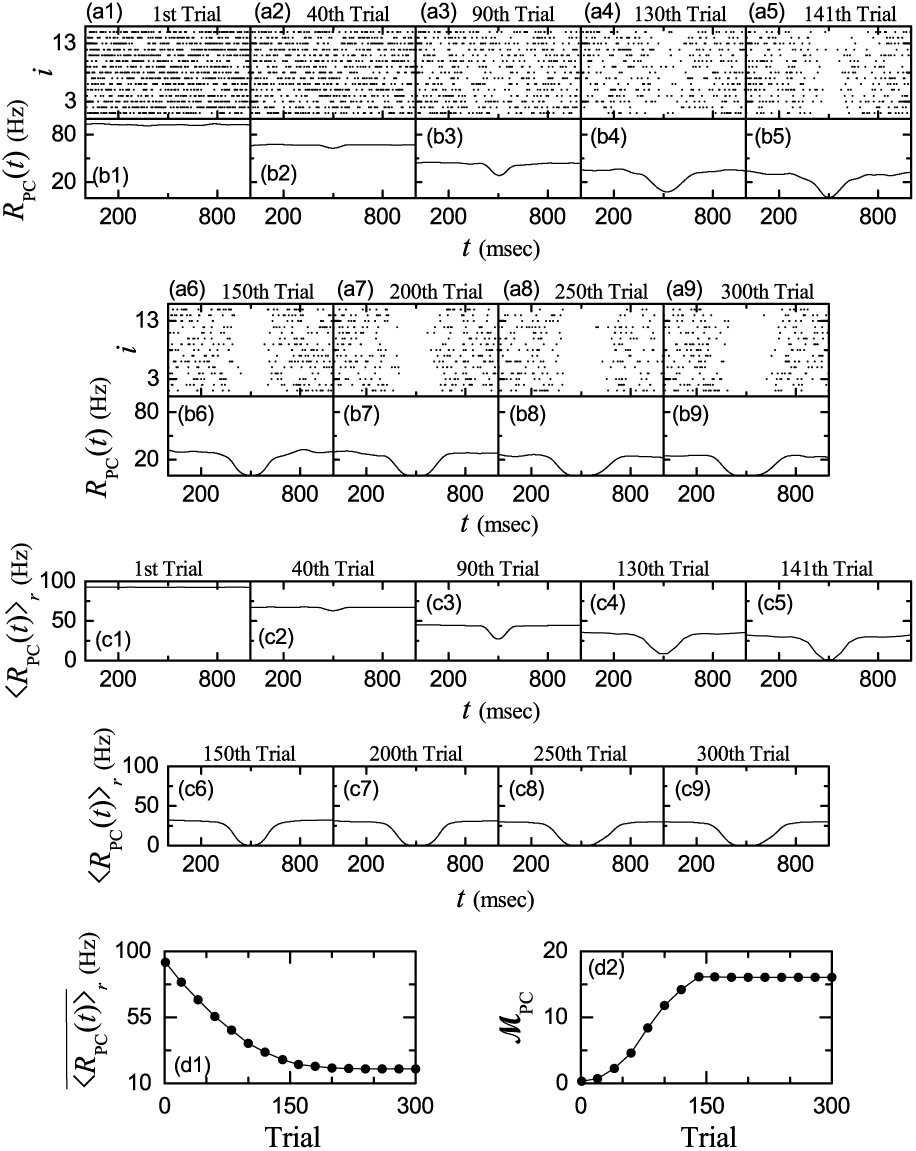
Change in firing activity of PCs during learning trial in the optimal case of 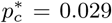. (a1)-(a9) Raster plots of spikes of PCs and (b1)-(b9) instantaneous population spike rates *R*_PC_(*t*). (c1)-(c9) Realization-averaged instantaneous population spike rates *(R*_PC_(*t*))_*r*_ ; number of realizations is 100. Plots of (d1) trial-averaged mean 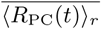 and (d2) modulations ℳ_PC_ for ⟨*R*_PC_(*t*)⟩_*r*_ versus trial.

As shown in Figs. 8(b1)-8(b9), bin-averaged normalized synaptic weights 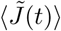 form a step-well-shaped curve. In the middle part of each trial, strong LTD occurs due to contribution of only well-matched firing group. On the other hand, at the initial and the final parts, somewhat less LTD takes place because both the ill-matched firing group (with practically no LTD) and the well-matched firing group make contributions together. As a result of this kind of effective depression at the (excitatory) PF-PC synapses, with increasing the learning trial, raster plots of spikes of all the 16 PCs become more and more sparse in the middle part of each trial (i.e, near *t* = 500 msec), which may be clearly seen in the instantaneous population spike rate ⟨*R*_PC_(*t*) ⟩_*r*_. ⟨*R*_PC_(*t*) ⟩_*r*_ becomes lower in the middle part than at the initial and the final parts. Thus, ⟨*R*_PC_(*t*) ⟨_*r*_ also forms a step-well-shaped curve with a minimum in the middle part.

As the trial is increased, such step-well-shaped curve for ⟨*R*_PC_(*t*) ⟩_*r*_ comes down and the (top) width and the depth of the well increase. Eventually, at the 141th trial, a “zero-bottom” is formed in the step-well in the middle part of the trial (i.e., ⟨*R*_PC_(*t*) ⟨ ≃_*r*_ 0 for 468 < *t* < 532 msec). Appearance of the zero-bottom in the step-well is the prerequisite condition for acquisition of CR. At the zero-bottom of the step-well, PCs stop inhibition completely. This process may be seen well in Figs. 10(c1)-10(c5). Thus, from the 141th threshold trial, the CN neuron may fire spikes which evoke CR, which will be seen in Fig. 11. With increasing the trial from the 141th trial, both the (top) width of the step-well and the zero-bottom width are increased, although the depth of the well remains unchanged [see Figs. 10(c6)-10(c9)]. As a result, the strength 𝒮of CR increases, while its timing degree 𝒯_*d*_ is decreased; the details will be given in Fig. 11. The (overall) learning efficiency degree ℒ_*e*_, taking into consideration both 𝒯_*d*_ and 𝒮, increases with the trial, and becomes saturated at about the 250th trial.

**FIG. 11:**
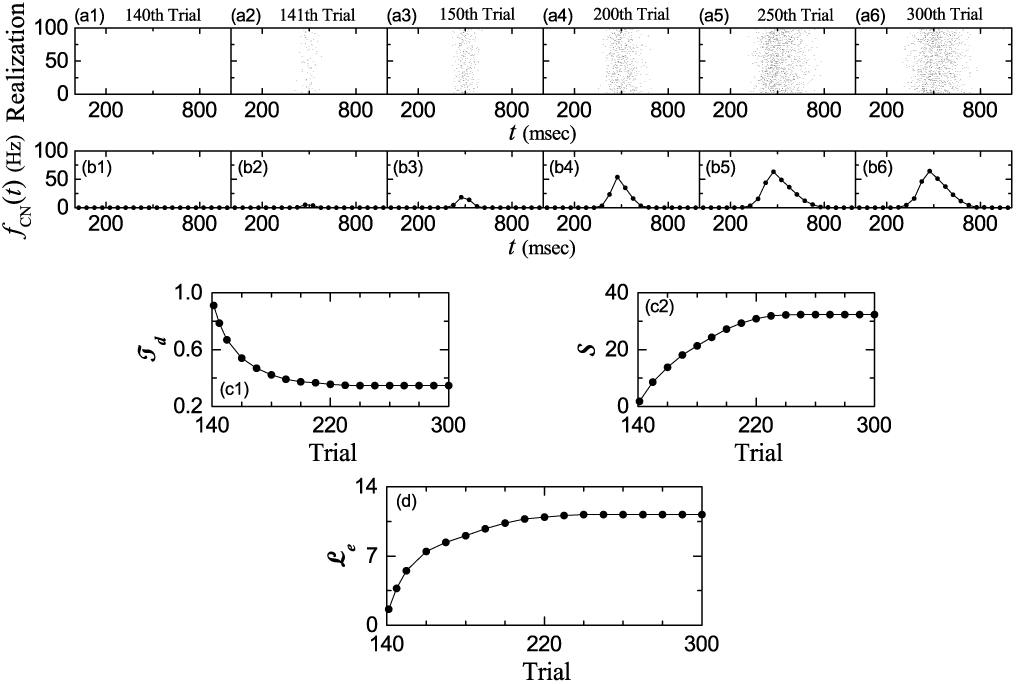
Change in firing activity of the CN neuron during learning trial in the optimal case of 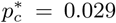. (a1)-(a6) Raster plots of spikes of the CN neuron (i.e., collection of spike trains for all the realizations; number of realizations is 100) and (b1)-(b6) bin-averaged instantaneous individual firing rate *f*_CN_(*t*); the bin size is Δ*t* = 50 msec. Plots of (c1) timing degree 𝒯_*d*_ and (c2) strength of CR versus trial. (d) Plot of (overall) learning efficiency degree ℒ_*e*_ for the CR versus trial.

Figures 10(d1) and 10(d2) show plots of trial-averaged mean 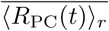 (i.e., time average of ⟨ (*R*_PC_(*t*)) ⟩_*r*_ over the trial) and modulation ℳPC of ⟨*R*_*PC*_(*t*)⟩_*r*_ versus trial, respectively. Due to effective LTD at the PF-PC synapses, the trial-averaged mean decreases 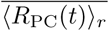 monotonically from 92.47 Hz, and it gets saturated at 19.91 Hz at about the 250th cycle. On the other hand, the modulation ℳPC increases monotonically from 0.352 Hz, and it becomes saturated at 16.11 Hz at about the 141th cycle. After the 141th threshold trial, ℳPC remains unchanged due to no change in the depth of the step-well, unlike the case of 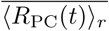. These PCs (principal cells of the cerebellar cortex) exert effective inhibitory coordination on the CN neuron which evokes the CR (i.e., learned eyeblink).

Figure 11 shows change in firing activity of the CN neuron which produces the final output of the cerebellum during learning trial in the optimal case of 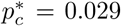. Trial-evolutions of raster plots of spikes of the CN neuron (i.e., collection of spike trains for all the realizations; number of realizations is 100) and the bin-averaged instantaneous individual firing rates *f*_CN_(*t*) (i.e., the number of spikes of the CN neuron in a bin with the bin width Δ*t* = 50 msec) are shown in Figs. 11(a1)-11(a6) and Figs. 11(b1)-11(b6), respectively. At the 140th trial, the CN neuron does not fire due to strong inhibition from the PCs, and thus it is silent (i.e., it lies in the silent period) during the whole trial stage (0 < *t* < 1000 msec). However, as a result of appearance of the zero-bottom in the step-well for ⟨*R*_*PC*_(*t*)⟩_*r*_ at the 141th threshold trial, the CN neuron begins to fire spikes in the middle part of the trial. In this case, as the time is increased from *t* = 0, the CN neuron first lies in the silent period, then a transition to the firing state occurs in the middle part, and finally another transition to the silent state also takes place. With increasing the trial, raster plots of spikes of the CN neuron become more and more dense in the middle part of each trial, in contrast to the case of PCs.

This process may be clearly seen in the instantaneous individual firing rates *f*_CN_(*t*). Due to the effective inhibitory coordinations of PCs on the CN neuron, *f*_CN_(*t*) begins to increase from 0 in the middle part of the trial, it reaches a peak, and then it decreases to 0 relatively slowly. The peak of *f*_CN_(*t*) appears a little earlier than the US presentation (*t* = 500 msec) which may denote the anticipatory CR [10, 49]. Thus, *f*_CN_(*t*) forms a bell-shaped curve. As the trial is increased, the bottom-base width and the peak height of the bell are increased, and *f*_CN_(*t*) seems to be saturated at about the 250th trial.

Figures 11(c1) and 11(c2) show plots of the timing degree 𝒯_*d*_ and the strength 𝒮 of the CR versus trial, respectively. The timing degree 𝒯_*d*_, representing the matching degree between the firing activity of the CN neuron [*f*_CN_(*t*)] and the US signal *f*_US_(*t*), is given by the cross-correlation *Corr*_T_(0) at the zero lag between *f*_CN_(*t*) and *f*_US_(*t*):

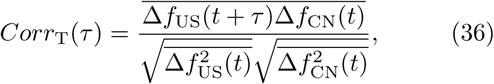

where 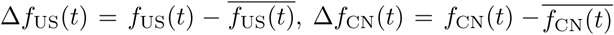, and the overline denotes the time average. Practically, 𝒯_*d*_ reflects the width of the bottom base of the bell curve. With increasing the trial, the width of the bottom base increases, due to increase in the (top) width of the step-well curve for the PCs. As a result, 𝒯_*d*_ decreases monotonically from 0.912 at the 141th trial, and it becomes saturated at 0.346 at about the 250th trial. On the other hand, as the trial is increased, the peak height of the bell increases. Thus, the strength 𝒮 of CR (representing the amplitude of eyelid closure), given by the modulation [(maximum - minimum)/2] of *f*_CN_(*t*), increases monotonically from 1.803 at the 141th trial, and it gets saturated at 32.38 at about the 250th trial.

Then, the (overall) learning efficiency degree *ℒ*_*e*_ for the CR, taking into consideration both the timing degree 𝒯 _*d*_ and the strength 𝒮 of CR, is given by their product:

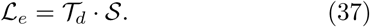

Figure 11(d) shows a plot of ℒ _*e*_ versus trial. ℒ _*e*_ increases monotonically from 1.645 at the 141th trial, and it becomes saturated at about the 250th cycle. Thus, we get the saturated learning efficiency degree 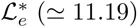. As will be seen in the next subsection, 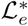 is the largest one among the available ones. Hence, in the optimal case of 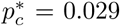 where firing patterns of GR clusters with the variety degree *𝒱* ^*^ (*≃* 1.842) are the most various, motor learning for the EBC with the saturated learning efficiency degree 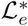 is the most effective.

Learning progress can be clearly seen in the IO system. During the learning trial, the IO neuron receives both the excitatory (airpuff) US signal for the desired timing and the inhibitory signal from the CN neuron (representing a realized eye-movement). Then, the learning progress degree *ℒ*_*p*_ is given by the ratio of the time-averaged inhibitory input from the CN neuron to the magnitude of the time-averaged excitatory input of the desired US timing signal:

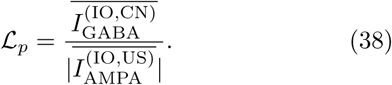

Here, 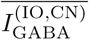 is the trial-averaged inhibitory GABA receptor-mediated current from the CN to the IO, and 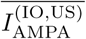 is the trial-averaged excitatory AMPA receptor-mediated current into the IO via the desired US timing signal; no (excitatory) NMDA receptors exist on the IO.[Note that the 4th term in Eq. (1) is given by *–*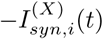, because 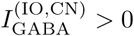 and 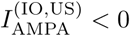.

Figure 12(a1) shows plots of 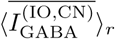 (open circles) and 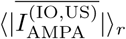 (crosses) versus trial in the optimal case of 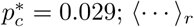 represents an average over 100 realizations. At the 141th threshold trial, acquisition of CR starts (i.e., the CN neuron begins to fire spikes). Hence, before the threshold the trial-averaged inhibitory input from the CN neuron is zero, it begins to increase from the threshold and converges to the constant trial-averaged excitatory input through the US signal for the desired timing. Thus, with increasing the trial, ℒ_*p*_ is zero before the threshold, it begins to increase from the threshold, and becomes saturated at ℒ_*p*_ = 1, as shown well in Fig. 12(a2). In this saturated case, the trial-averaged excitatory and inhibitory inputs to the IO are balanced.

**FIG. 12:**
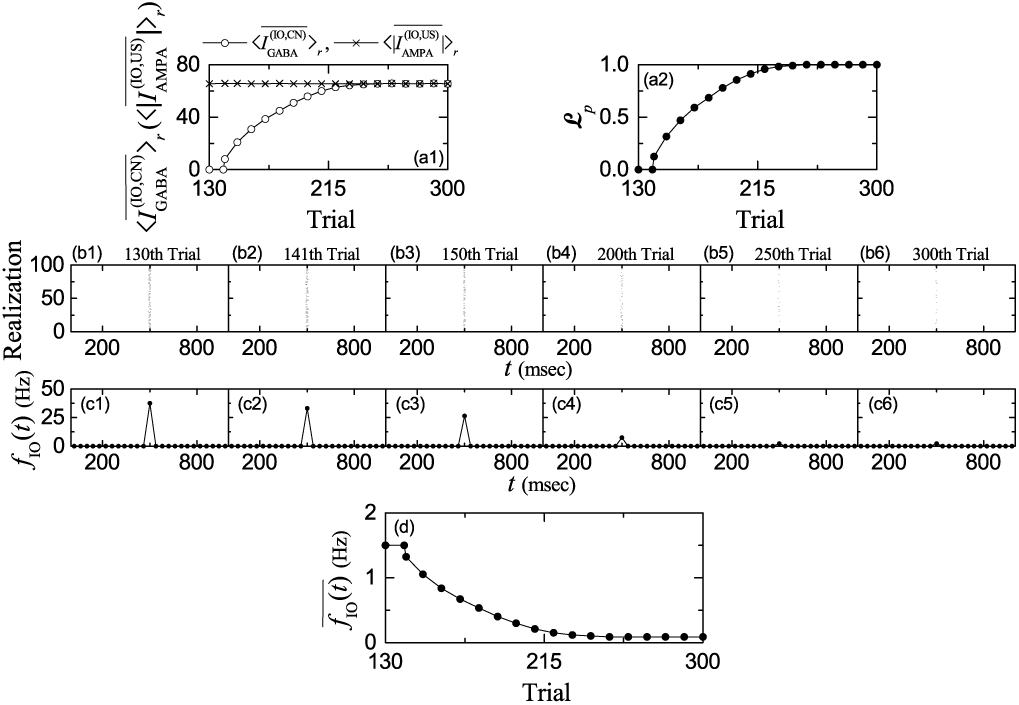
Change in firing activity of the IO neuron during learning trial in the optimal case of 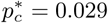. Plots of (a1) realization-average for the time<u>-</u>averaged inhibitory synaptic current from the CN neuron 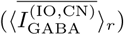 (open circles) and realization-average for the time-averaged excitatory synaptic current via the (airpuff) US signal 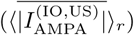 versus trial; number of realizations ⟨…⟩_*r*_ is 100. (a2) Plot of learning progress degree ℒ_p_versus trial. (b1)-(b5) Raster plots of spikes of the IO neuron (i.e., collection of spike trains for all the realizations; number of realizations is 100) and (c1)-(c5) bin-averaged instantaneous individual firing rate *f*_IO_(*t*); the bin size is Δ*t* = 40 msec. (d) Plot of trial-averaged individual firing rate 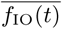 versus trial.

We also investigate the firing activity of IO neuron during learning process. Figures 12(b1)-12(b6) and Figures 12(c1)-12(c6) show trial-evolutions of raster plots of spikes of the IO neuron (i.e., collection of spike trains for all the realizations; number of realizations is 100) and the bin-averaged instantaneous individual firing rates *f*_IO_ (i.e., the number of spikes of the IO neuron in a bin with the bin width Δ*t* = 40 msec), respectively. Before the 141th threshold trial, relatively dense spikes appear in the middle part of the trial in the raster plot of spikes, due to the effect of excitatory US signal. However, with increasing the trial from the threshold, spikes in the middle part become sparse, because of increased inhibitory input from the CN neuron. In this case, the bin-averaged instantaneous individual firing rate *f*_IO_(*t*) of the IO neuron forms a bell-shaped curve due to the US signal into the IO neuron. With increasing the trial from the 141th threshold, the amplitude of *f*_IO_(*t*) begins to decrease due to the increased inhibitory input from the CN neuron, and it becomes saturated at about the 250th trial. Thus, the trial-averaged individual firing rate 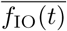 is constant (=1.5 Hz) before the threshold without the inhibitory input from the CN neuron. However, with increasing the trial from the threshold, it is decreased from 1.326 Hz due to increase in the inhibitory input from the CN neuron, and gets saturated at 0.0902 Hz at about the 250th trial, as shown in Fig. 12(d).

The firing output of the IO neuron is fed into the PCs via the CFs. Hence, with increasing the trial from the threshold, the error-teaching CF instructor signals become weaker and saturated at about the 250th cycle. While the saturated CF signals are fed into the PCs, trial-level balance between ΔLTD and ΔLTP occurs at the PF-PC synapses, as shown in Figs. 9(c1) and 9(c2). As a result, saturation for the trial-averaged bin-averaged synaptic weights 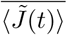 appears [see Fig. 8(c)]. Then, the subsequent learning process in the PC-CN system also gets saturated, and we obtain the saturated learning efficiency degree 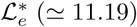, which is shown in Fig. 11(d).

### E. Variation of The Connection Probability *p*_*c*_ and Strong Correlation between Variety Degree *𝒱* and Efficiency Degree *ℒ*_*e*_ of CR

In the above subsections, we consider only the optimal case of 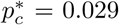 (i.e., 2.9%) where the firing patterns of the GR clusters are the most various. From now on, we change the connection probability *p*_*c*_ from GO to GR cells, and study dependence of the variety degree *𝒱* for the firing patterns in the GR clusters and the learning efficiency degree *ℒ*_*e*_ of CR on *p*_*c*_.

We first consider both the highly-connected case of *p*_*c*_ = 0.3 (i.e., 30%) and the lowly-connected case of *p*_*c*_ = 0.003 (i.e., 0.3%). Figures 13(a1) and 13(a2) show the raster plots of spikes of 10^3^ randomly chosen GR cells for *p*_*c*_ = 0.3 and 0.003, respectively. The population-averaged firing activities in the whole population of GR cells may be well seen in the instantaneous whole-population spike rates *R*_GR_(*t*) in Figs. 13(b1) and 13(b2) for *p*_*c*_ = 0.3 and 0.003, respectively.

**FIG. 13:**
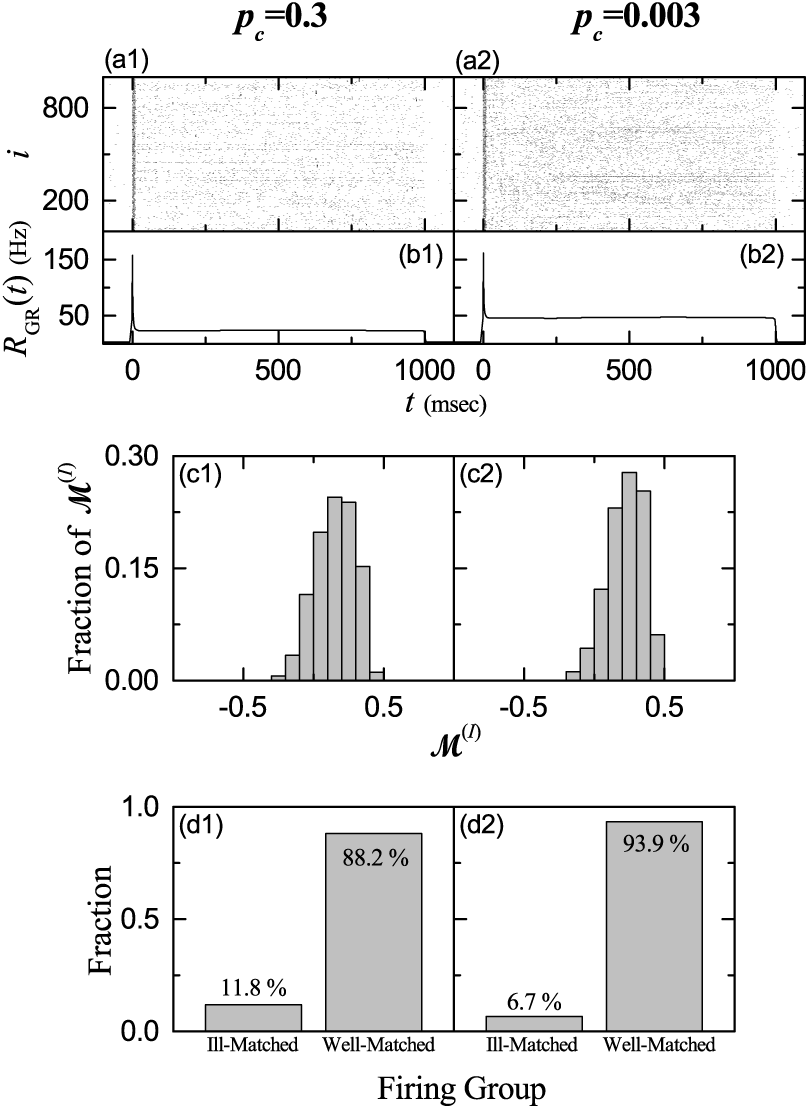
Highly-connected (*p*_*c*_ = 0.3) and lowly-connected (*p*_*c*_ = 0.003) cases. Raster plots of spikes of 10^3^ randomly chosen GR cells for (a1) *p*_*c*_ = 0.3 and (a2) *p*_*c*_ = 0.003. Instantaneous whole-population spike rates *R*_GR_(*t*) in the whole population of GR cells for (b1) *p*_*c*_ = 0.3 and (b2) *p*_*c*_ = 0.003. Band width for *R*_GR_(*t*): *h* = 10 msec. Distributions of matching indices {ℳ^(*I*)^} of the firing patterns in the GR clusters in the whole population for (c1) *p*_*c*_ = 0.3 and (c2) *p*_*c*_ = 0.003. Bin size for the histograms in (c1) and (c2) is 0.1. Fractions of well-matched and ill-matched firing groups for (d1) *p*_*c*_ = 0.3 and (d2) *p*_*c*_ = 0.003.

As shown in Fig. 2(b), each GR cluster is bounded by four glomeruli (corresponding to the terminals of the MFs) at both ends. Each glomerulus receives inhibitory inputs from nearby 81 GO cells with the connection probability *p*_*c*_. In the highly-connected case of *p*_*c*_ = 0.3, on average, about 24 GO cell axons innervate each glomerulus. Then, each GR cell in a GR cluster receives about 97 inhibitory inputs via four dendrites which contact the four glomeruli at both ends. In this highly-connected case, inhibitory inputs from the pre-synaptic GO cells are increased, in comparison with the optimal case of 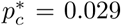. As a result, spike density in the raster plot of spikes is decreased (i.e., spikes become sparse) due to decreased individual firing rates, and hence the flat top part of *R*_GR_(*t*) becomes lowered, in comparison to the optimal case in Fig. 4(b).

In the highly-connected case of *p*_*c*_ = 0.3, differences between total inhibitory synaptic inputs from pre-synaptic GO cells to each GR cells are decreased due to increase in the number of pre-synaptic GO cells. In addition, the excitatory inputs into each GR cells via MFs are Poisson spike trains with the same firing rates, and hence they are essentially the same. Hence, differences between the total synaptic inputs (including both the inhibitory and the excitatory inputs) into each GR cells become reduced. These less different inputs into GR cells produce less different outputs (i.e. firing activities) of GR cells, which become more similar to the population-averaged firing activity *R*_GR_(*t*) with a flat top in Fig. 13(b1). Thus, GR cells tend to exhibit relatively regular firings during the whole trial stage (0 < *t* < 1000 msec), in comparison with optimal case of 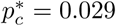. Consequently, the raster plot of sparse spikes for *p*_*c*_ = 0.3 becomes relatively uniform [compare Fig. 13(a1) with Fig. 4(a)].

On the other hand, in the lowly-connected case of *p*_*c*_ = 0.003, the inhibitory inputs from GO cells into GR cells are so much reduced, and the excitatory MF signals into the GR cells become dominant inputs. Hence, spike density in the raster plot of spikes is increased (i.e., spikes become dense), because of increased individual firing rate, and the flat top part of *R*_GR_(*t*) becomes raised, in comparison to the optimal case in Fig. 4(b). Furthermore, differences between the total synaptic inputs into each GR cells become reduced, because the dominant excitatory MF signals, generated by the Poisson spike trains with the same firing rates, are essentially the same, and thus firing activities of GR cells become more similar to *R*_GR_(*t*) with a flat top in Fig. 13(b2). Hence, as in the highly-connected case, GR cells tend to show relatively regular firings during the whole trial stage. As a result, the raster plot of dense spikes for *p*_*c*_ = 0.003 also becomes relatively uniform, as shown in Fig. 13(a2), in comparison with the optimal case in Fig. 4(a).

Figures 13(c1) and 13(c2) show the distributions of matching indices {ℳ^(*I*)^}for *p*_*c*_ = 0.3 and 0.003, respectively. The ranges in the distributions of {ℳ^(*I*)^} for *p*_*c*_ = 0.3 and 0.003 are (−0.21, 0.44) and (−0.18, 0.48), respectively. In both cases, their ranges are narrowed from both the positive and the negative sides, in comparison with the range (−0.49, 0.79) in the optimal case of 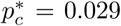. As explained above, in both the highly- and the lowly-connected cases of *p*_*c*_ = 0.3 and 0.003, GR cells tend to exhibit relatively regular firings in the whole trial stage, due to decrease in the differences in the total synaptic inputs from GO cells into each GR cells, which is in contrast to the optimal case of 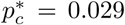 where random repetitions of transitions between bursting and silent states (both of which are persistent long-lasting ones) occur. Then, in both the highly- and the lowly-connected cases, highly well-matched firing patterns with higher ℳ^(*I*)^ and highly ill-matched firing patterns with higher magnitude |ℳ^(*I*)^ | disappear, which leads to reduction in the ranges of the distributions of {ℳ^(*I*)^} arise.

Due to the narrowed distribution of {ℳ^(*I*)^}, both the mean (≃0.239) and the standard deviation (≃0.360) in the highly-connected case of *p*_*c*_ = 0.3 are decreased, in comparison to the optimal case where the mean and the standard deviation are 0.333 and 0.614, respectively. Then, the variety degree for the firing patterns 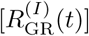 in the GR clusters, denoting a quantitative measure for various recoding in the granular layer, is given by the relative standard deviation for the distribution of {ℳ^(*I*)^} [see Eq. (27)]. For *p*_*c*_ = 0.3, its variety degree is *𝒱*≃ 1.506 which is smaller than *𝒱*^*^ (≃ 1.842) in the optimal case of 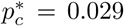. Similar to the highly-connected case, for *p*_*c*_ = 0.003 both the mean (≃ 0.272) and the standard deviation (≃ 0.315) for the distribution of {ℳ^(*I*)^} are also decreased. In this lowly-connected case, the variety degree *𝒱* for the firing patterns 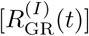 in the GR clusters is *𝒱* ≃1.157 which is much smaller than *𝒱*^*^ (≃ 1.842) in the optimal case. We also note that the variety degree 𝒱 for *p*_*c*_ = 0.003 is smaller than that for *p*_*c*_ = 0.3; *𝒱*^*^ = 1.842 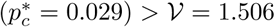 (*p*_*c*_ = 0.3) > *𝒱* = 1.157 (*p*_*c*_ = 0.003).

Figures 13(d1) and 13(d2) show fractions of well-matched ({ℳ^(*I*)^ > 0}) and ill-matched ({ℳ^(*I*)^ > 0}) firing groups for *p*_*c*_ = 0.3 and 0.003, respectively. In the highly-connected case of *p*_*c*_ = 0.3, the well-matched firing group is a major one with fraction 88.2%, while the ill-matched firing group is a minor one with fraction 11.8%. In comparison with the optimal case where the fraction of well-matched firing group is 82.1%, the fraction of well-matched firing group for *p*_*c*_ = 0.3 is increased. In this highly-connected case, the firing-group ratio, given by the ratio of the fraction of the well-matched firing group to that of the ill-matched firing group, is ℛ_sp_ ≃ 7.48 which is larger than that 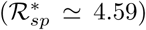 in the optimal case. In the lowly-connected case of *p*_*c*_ = 0.003, the fraction of well-matched firing group is more increased to 93.9%. Hence, the firing-group ratio is ℛ_sp_ 13.8 which is much larger than that in the optimal case.

Due to decrease in differences between the total synaptic inputs into each GR cells, firing activities of GR cells for *p*_*c*_ = 0.3 and 0.003 become more similar to the population-averaged firing activity *R*_GR_(*t*), in comparison with the optimal case of 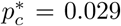. We note that *R*_GR_(*t*) is well-matched with the US signal *f*_US_(*t*) [i.e, *R*_GR_(*t*) has a positive conjunction index with respect to *f*_US_(*t*)], which results in increase in the fraction of well-matched firing group for *p*_*c*_ = 0.3 and 0.003. In contrast to the case of *p*_*c*_ = 0.3, in the lowly-connected case of *p*_*c*_ = 0.003, inhibitory inputs into each GR cells are so much reduced, and hence the dominant inputs are just the excitatory MF signals which are well-matched with the US signal *f*_US_(*t*). Thus, the fraction of well-matched firing group for *p*_*c*_ = 0.003 becomes larger than that for *p*_*c*_ = 0.3.

These changes in the variety degree *𝒱* of the firing patterns in the GR clusters have direct effect on the synaptic plasticity at the PF-PC synapses and the subsequent learning process in the PC-CN system. As shown in the optimal case of 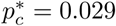, the ill- and the well-matched firing groups play their own roles for the CR. The ill-matched firing group plays a role of protection barrier for the timing of CR, while the strength of CR is determined by strong LTD in the well-matched firing group. Due to break-up of highly well-matched and highly ill-matched firing patterns, the distributions of {ℳ^(*I*)^} for the highly- and the lowly-connected cases of *p*_*c*_ = 0.3 and 0.003 are narrowed. Hence, both the timing degree 𝒯_*d*_ and the strength 𝒮 of CR are decreased for both *p*_*c*_ = 0.3 and 0.003, which are well shown in Figs. 14 and 15.

**FIG. 14:**
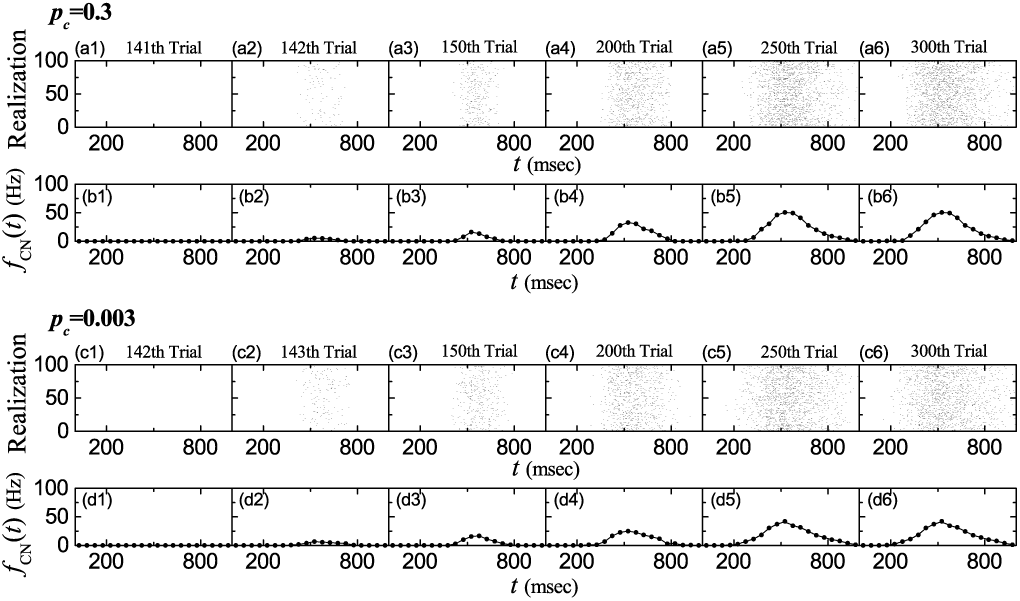
Change in firing activity of the CN neuron in the highly-connected (*p*_*c*_ = 0.3) and the lowly-connected (*p*_*c*_ = 0.003) cases. Case of *p*_*c*_ = 0.3: (a1)-(a6) raster plots of spikes of the CN neuron (i.e., collection of spike trains for all the realizations; number of realizations is 100) and (b1)-(b6) bin-averaged instantaneous individual firing rate *f*_CN_(*t*); the bin size is Δ*t* = 50 msec. Case of *p*_*c*_ = 0.003: (c1)-(c6) raster plots of spikes of the CN neuron and (d1)-(d6) bin-averaged instantaneous individual firing rate *f*_CN_(*t*).

**FIG. 15:**
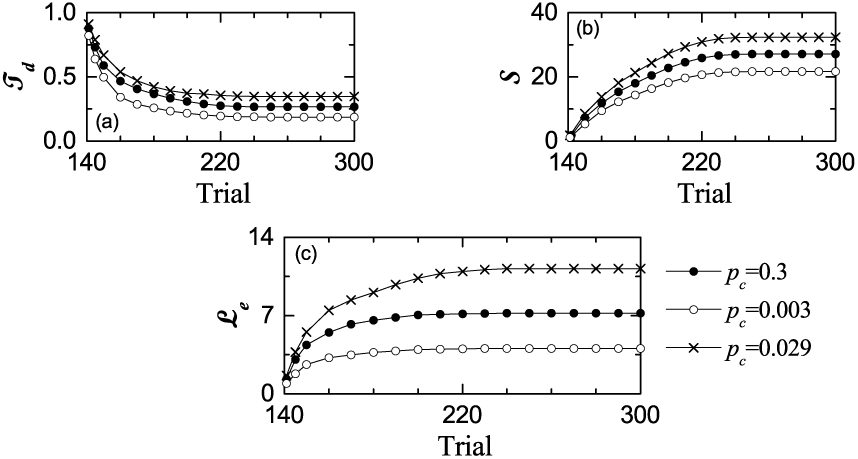
Timing degree, strength, and learning efficiency degree of the CR in the highly-connected (*p*_*c*_ = 0.3) and the lowly-connected (*p*_*c*_ = 0.003) cases. (a) Plots of timing degree 𝒯_*d*_ of the CR versus trial. (b) Plots of strengths 𝒮of the CR versus trial. (c) Plots of learning efficiency degree ℒ_*e*_ for the CR versus trial. Solid circles, open circles, and crosses represent data in the cases of *p*_*c*_ = 0.3, 0.003, and 0.029, respectively.

**FIG. 16:**
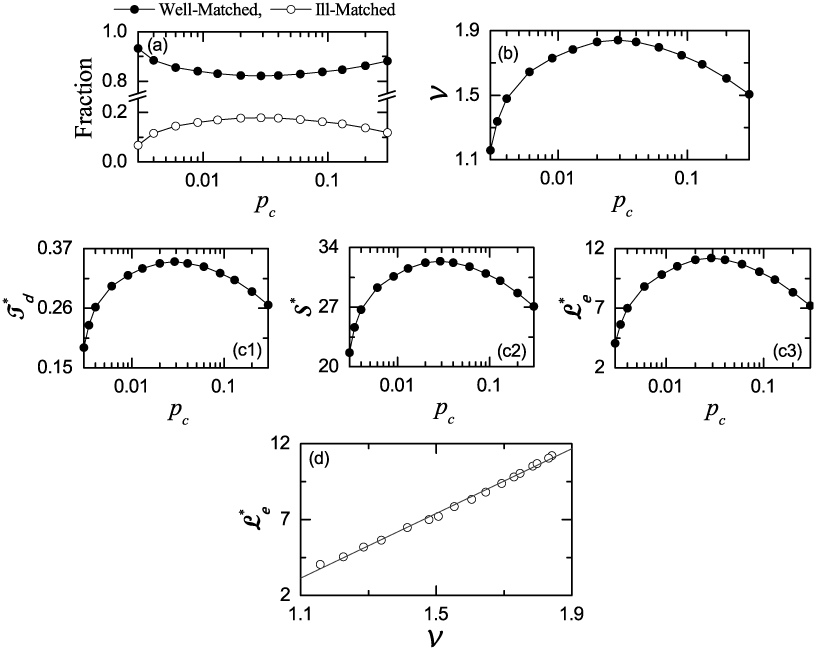
Strong correlation between the variety degree *𝒱* and the saturated learning efficiency degree _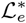_ (a) Fractions of well-matched (solid circles) and ill-matched (open circles) firing groups versus the connection probability *p*_*c*_. (b) Plot of variety degree *𝒱* of the firing patterns in the GR clusters versus *p*_*c*_. Plots of saturated (c1) timing degree 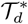, (c2) strengths 𝒮^*^, and (c3) learning efficiency degree 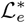 of the CR versus *p*_*c*_. (d) Plot of 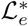 versus *𝒱*

Figure 14 shows change in the firing activity of the CN neuron which generates the final output of the cerebellum during learning trial in the highly- and the lowly-connected cases of *p*_*c*_ = 0.3 and 0.003. For *p*_*c*_ = 0.3, trial-evolutions of the raster plots of spikes of the CN neuron (i.e., collection of spike trains for all the realizations; number of realizations is 100) and the bin-averaged instantaneous individual firing rates *f*_CN_(*t*) (i.e., the number of spikes of the CN neuron in a bin with the bin width Δ*t* = 50 msec) are shown in Figs. 14(a1)-14(a6) and Figs. 14(b1)-14(b6), respectively. In this highly-connected case, at the 142th threshold trial, the CN neuron begins to fire in the middle part of the trial. Thus, acquisition of CR occurs a little later, in comparison with the optimal case of 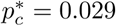 with the 141th threshold. In this case, *f*_CN_(*t*) forms a bell-shaped curve. With increasing the trial from the threshold, raster plots of spikes of the CN neuron become more and more dense in the middle part of each trial, and bottom-base width and peak height of the bell curve for *f*_CN_(*t*) increase. At about the 250th trial, *f*_CN_(*t*) seems to become saturated.

For *p*_*c*_ = 0.3, the bottom-base width of the bell curve is wider and its peak height is shorter, in comparison with the optimal case (see Fig. 11). Due to break-up of highly ill-matched firing patterns (which play the role of protection barrier for timing of CR), bottom-base width (associated with the reciprocal of timing degree of CR) of the bell increases. Also, peak height of the bell (related to the strength of CR) decreases because of break-up of highly well-matched firing patterns (which induce strong LTD and determine the strength of CR). Consequently, as the variety degree *𝒱* of the firing patterns in the GR clusters is deceased from 1.842 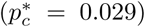 to 1.507 (*p*_*c*_ = 0.3), the bottom-base width of the bell curve is increased, and the peak height is decreased.

For *p*_*c*_ = 0.003, trial-evolutions of the raster plots of spikes of the CN neuron and the bin-averaged instantaneous individual firing rates *f*_CN_(*t*) are shown in Figs. 14(c1)-14(c6) and Figs. 14(d1)-14(d6), respectively. In this lowly-connected case, acquisition of CR occurs at the 143th trial which is a little later in comparison with the 142th threshold for *p*_*c*_ = 0.3. Similar to the highly-connected case, with increasing the trial from the threshold, raster plots of spikes of the CN neuron become more and more dense in the middle part of each trial, and the bottom-base width and the peak height of the bell curve for *f*_CN_(*t*) increase. Eventually, saturation occurs at about the 250th trial. In comparison to the highly-connected case, the bottom-base width of the bell curve is wider and its peak height is shorter, because of more break-up of highly ill-matched firing patterns and highly well-matched firing patterns (which results in more decrease in the variety degree of firing patterns). As a result, with decreasing the variety degree 𝒱 from 1.507 (*p*_*c*_ = 0.3) to 1.157 (*p*_*c*_ = 0.003), the bottom-base width of the bell curve increases, and the peak height decreases (i.e., less variety in the firing patterns results in decrease in the timing degree and the strength of CR).

Figures 15(a) and 15(b) show plots of the timing degree 𝒾_*d*_ and the strength of CR versus trial for *p*_*c*_ = 0.3 (solid circles), 0.003 (open circles), and 0.029 (crosses), respectively. The timing degree 𝒾_*d*_, denoting the matching degree between the firing activity of the CN neuron [*f*_CN_(*t*)] and the US signal *f*_US_(*t*) for a timing, is given by the cross-correlation *Corr*_T_(0) at the zero lag between *f*_CN_(*t*) and *f*_US_(*t*) in Eq. (36). This timing degree 𝒾_*d*_ reflects the width of the bottom base of the bell curve. With increasing the trial, the width of the bottom base increases, as shown in Fig. 14, and hence 𝒾_*d*_ decreases monotonically, and it becomes saturated at about the 250th trial. We note that, as the variety degree 𝒱 of the firing patterns is decreased (*𝒱* = 1.507, 1.157, and 1.842 for *p*_*c*_ = 0.3, 0.003, and 0.029, respectively), 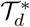 (saturated value of *𝒾*_*d*_) decreases; 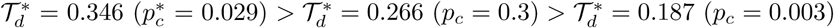.

On the other hand, as the trial is increased, the peak height of the bell increases. Hence, the strength 𝒮of CR, given by the modulation of *f*_CN_(*t*), increases monotonically, and it becomes saturated at about the 250th trial. In this case, 𝒮^*^ (saturated value of) also is decreased with decrease in the variety degree ; 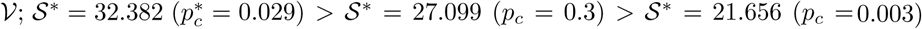.

We then consider the learning efficiency degree *ℒ*_*e*_ for the CR, given by product of the timing degree 𝒯_*d*_ and the strength 𝒮in Eq. (37). Figure 15(c) shows a plot of ℒ_*e*_ versus trial. ℒ_*e*_ increases monotonically from the threshold trial, and it becomes saturated at about the 250th cycle. Thus, we obtain the saturated learning efficiency degree ^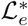^, the values of which are 7.216, 4.054, and 11.19 for *p*_*c*_ = 0.3, 0.003, and 0.029, respectively. Among the three cases, 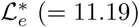 in the optimal case is the largest one, and 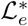 is decreased with decrease in the variety degree 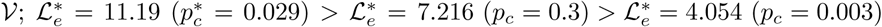.

Finally, based on the above two examples for the highly- and the lowly-connected cases, we investigate dependence of the variety degree 𝒱 for the firing patterns of the GR clusters and the saturated learning efficiency degree 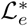on *p*_*c*_ by varying it from the optimal value 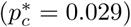. Figure 16(a) shows plots of fractions of the well- and the ill-matched firing groups versus *p*_*c*_. The fraction of the well-matched firing group (solid circles) forms a well-shaped curve with a minimum in the optimal case of 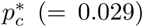, while the fraction of the ill-matched firing group (open circles) forms a bell-shaped curve with a maximum in the optimal case. In the optimal case of 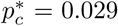, the firing-group ratio (i.e., ratio of fraction of the well-matched firing group to fraction of the ill-matched firing group) is 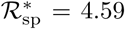. As *p*_*c*_ is changed (i.e., increased or decreased) from 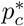, the fraction of the well-matched firing group increases, and then the firing-group ratio ℛ _sp_ increases from ^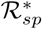^ in the optimal case.

Figure 16(b) show a plot of the variety degree *𝒱* for the firing patterns in the GR clusters. The variety degrees *𝒱* forms a bell-shaped curve with a maximum *𝒱*^*^ (≃1.842) in the optimal case of 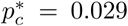. With changing *p*_*c*_ from 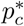, 𝒱 decreases from ^*^. Hence, in the optimal case, temporal recoding of GR cells is the most various. Figures 16(c1)-16(c3) show plots of the saturated timing degree 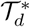, the saturated strength *𝒮*^*^, and the saturated learning efficiency degree 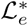 of CR, respectively. All of them form bell-shaped curves with maxima 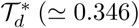, and 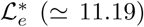 in the optimal case of 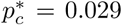. As *p*_*c*_ is changed from 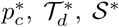, and 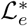 are decreased. Figure 16(d) shows a plot of 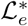 versus _*𝒱*_. As shown clearly in Fig. 16(d), both ^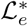^ and 𝒱 have a strong correlation with the Pearson’s correlation coefficient *r* ≃0.9982. Consequently, the more various in temporal recoding of the GR cells, the more effective in learning for the Pavlovian EBC.

## IV. SUMMARY AND DISCUSSION

We are concerned about the Pavlovian EBC. Various works on the EBC have been done experimentally in many mammalian species such as humans, monkeys, dogs, ferrets, rabbits, rats, and mice [14–18, 24–27]. Also, computational works reproduced some features (e.g., representation of time passage) of the EBC in artificial models [39–44], a realistic biological model [45–47], a rate-coding model [48], and a spiking neural network model [49]. However, more clarification is necessary for influences of various temporal recoding in GR clusters on the Pavlovian EBC.

To the best of our knowledge, for the first time, we made complete quantitative classification of various firing patterns in the GR clusters in terms of the newly-introduced matching index {ℳ^(*I*)^} and variety degree in the case of Pavlovian EBC. Each firing pattern is characterized in terms of its matching index ℳ^(*I*)^ between the firing pattern and the US signal for the desired timing. Then, the whole firing patterns are clearly decomposed into the well-matched (ℳ^(*I*)^ > 0) and the ill-matched (ℳ^(*I*)^ < 0) firing groups. Furthermore, the degree of various recoding of the GR cells may be quantified in terms of the variety degree, given by the relative standard deviation in the distribution of {ℳ^(*I*)^}. Thus, provides a quantitative measure for various temporal recoding of GR cells. It has also been shown that various total synaptic inputs (including both the excitatory inputs via MFs and the inhibitory inputs from the pre-synaptic GO cells) into the GR clusters result in generation of various firing patterns (i.e. outputs) in the GR clusters.

Based on the above dynamical classification of various firing patterns in the GR clusters, we made clear investigations on the influence of various recoding of GR cells on the Pavlovian EBC (i.e., their effect on the synaptic plasticity at the PF-PC synapses and the subsequent learning process in the PC-CN-IO system). To the best of our knowledge, this kind of approach, based on the well- and the ill-matched firing groups, is unique in studying the Pavlovian EBC. The well-matched firing patterns are strongly depressed by the error-teaching CF signals, and they make dominant contributions in the middle part of each trial. In contrast, for the ill-matched firing patterns with central gaps in the middle part, practically no LTD occurs due to no matching with the CF signals. Thus, in the middle part of each trial, strong LTD occurs via dominant contributions of well-matched firing group. On the other hand, at the initial and the final parts of each trial, less LTD takes place because both the ill-matched firing group with practically no LTD and the well-matched firing group make contributions together. As a result, a big modulation in bin-averaged synaptic weight 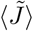 arises via interplay of the well- and ill-matched firing groups.

Due to this type of effective synaptic plasticity at the PF-PC synapses, the (realization-averaged) population spike rate ⟨*R*_PC_(*t*) ⟩ of PCs forms a step-well-shaped curve with a minimum in the middle part of each trial. When passing a threshold trial, a “zero-bottom” (where ⟨*R*_PC_(*t*) ⟩ ≃0) appears in the central well. At this threshold trial, the CN neuron begins to fire in the middle part of trial. Hence, appearance of the zero-bottom in the step-well for ⟨*R*_PC_(*t*) ⟩ is a prerequisite condition for acquisition of CR. In the subsequent trials, the individual firing rate *f*_CN_(*t*) of the CN neuron forms a bell-shaped curve with a maximum in the middle part (which is updown reversed with respect to ⟨*R*_PC_(*t*) ⟩). Outside the bottom of the bell, the CN neuron cannot fire, due to inhibition of ill-matched firing group (with practically no LTD). Hence, the ill-matched firing group plays a role of protection barrier for timing, and the timing degree of CR is reciprocally associated with the bottom width of the bell. In this case, the peak of the bell in the middle part is formed due to strong LTD in the well-matched firing group, and its height is directly related to the strength of CR (corresponding to the amplitude of eyelid closure). In this way, both the well- and the ill-matched firing groups play their own roles for the timing and the strength of CR, respectively.

By changing *p*_*c*_, we investigated dependence of the variety degree *𝒱* of the firing patterns and the saturated learning efficiency degree 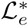 for the CR (given by the product of the timing degree and the strength of CR) on *p*_*c*_. Both 𝒱 and ^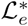^ have been found to form bell-shaped curves with peaks (𝒱^*^ ≃ 1.842 and _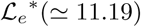_) at the same optimal value of 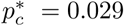. In Refs. [4, 49] where the parameter values were taken, based on physiological data, the average number of nearby GO cell axons innervating each GR cell is about 8, which is very close to that in the optimal case. Thus, we hypothesize that the granular layer in the cerebellar cortex has evolved toward the goal of the most various recoding. Moreover, Both 𝒱 and ^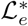^ have also been found to have a strong correlation with the Pearson’s correlation coefficient *r* ≃0.9982. Hence, the more various in the firing patterns of GR cells, the more efficient in learning for the Pavlovian EBC, which is the main result in our work.

To examine our main result, we also suggest a real experiment for the EBC. To control *p*_*c*_ in a given species of animals (e.g., a species of rabbit, rat, or mouse) in an experiment seems to be practically difficult, in contrast to the case of computational neuroscience where *p*_*c*_ may be easily varied. Instead, we may consider an experiment for several species of animals (e.g., 3 species of rabbit, rat, and mouse). In each species, a large number of randomly chosen GR cells (*i* = 1,, *M*) are considered. Then, through many CS-US trials, one may get the peristimulus time histogram (PSTH) for each *i*th GR cell [i.e., (bin-averaged) instantaneous individual firing rate 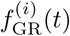 of the *i*th GR cell]. GR cells are expected to exhibit various PSTHs. Then, in the case of each *i*th GR cell, one can get its matching index ℳ_*i*_ between its PSTH 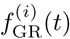 and the CF signal for the desired timing [i.e., the PSTH of the IO neuron *f*_IO_(*t*)]. In this case, the conjunction index *ℳ*_i_ is given by the cross-correlation at the zero-time lag between 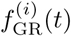 and *f*_IO_(*t*). Thus, one may get the variety degree of PSTHs of GR cells, given by the relative standard deviation in the distribution of {ℳ_*i*_ *}*, for the species.

In addition to the PSTHs of GR cells, under the many CS-US trials, one can also obtain a bell-shaped PSTH of a CN neuron [(bin-averaged) instantaneous individual firing rate *f*_CN_(*t*) of the CN neuron]. The reciprocal of bottom-base width and the peak height of the bell curve correspond to timing degree 𝒯_*d*_ and strength 𝒮for the EBC, respectively. In this case, the (overall) learning efficiency degree ℒ_*e*_ for the EBC is given by the product of 𝒯_*d*_ and. In this way, a set of (𝒱, *ℒ*_*e*_) may be experimentally obtained for each species, and depending on the species, the set of (𝒱, *ℒ*_*e*_) may change. Then, for example in the case of 3 species of rabbit, rat, and mouse, with the three different data sets for (𝒱, *ℒ*_*e*_), one can examine our main result (i.e., whether more variety in PSTHs of GR cells leads to more efficient learning for the EBC).

Finally, we make discussion on limitations of our present work and future works. In the present work where the ISI between the onsets of CS and US was set at 500 msec, we investigated the effect of various temporal recoding of GR cells on the Pavlovian EBC. The acquisition rate and the timing degree and strength of CR have been known to depend on the ISI [49, 85]. Hence, in a future work, it would be interesting to study dependence of EBC on the ISI. Based on the results of our work, it would also be interesting to study extinction of CR, as a future work. After acquisition of CR, we turn off the airpuff US. Then, the CR is expected to become gradually extinct via LTP at the PF-PC synapses [7]. In this work, we considered only the PF-PC synaptic plasticity. In the cerebellum, synaptic plasticity takes place at various synapses [86, 87] (e.g., MF-CN and PC-CN synapses [88], PF-BC and BC-PC synapses [89], and MF-GR cells synapses [90]). Hence, it would be interesting to make a future study on the influence of various synaptic plasticity at various synapses on the cerebellar learning for the Pavlovian EBC; particularly, we are interested in the effect of the synaptic plasticity at the MF-CN synapse on the EBC [7]. In addition to change in *p*_*c*_ (i.e., connection probability from GO to GR cells), one can vary synaptic inputs into the GR cells by changing NMDA receptor-mediated maximum conductances 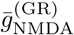and 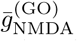[49]. Hence, as a future work, It would also be interesting to study the influence of NMDA receptor-mediated synaptic inputs on various recoding of GR cells and learning for the EBC by varying 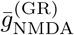 and 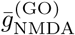.

## Acknowledgments

This research was supported by the Basic Science Research Program through the National Research Foundation of Korea (NRF) funded by the Ministry of Education (Grant No. 20162007688).

## Appendix: Parameter Values for The LIF Neuron Models and The Synaptic Currents

In this appendix, we list two tables which show parameter values for the LIF neuron models in Subsec. II C and the synaptic currents in Subsec. II D. These values are adopted from physiological data [4, 49].

For the LIF neuron models, the parameter values for the capacitance *C*_*X*_, the leakage current 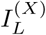, the AHP current 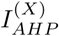, and the external constant current 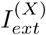 are shown in Table I.

For the synaptic currents, the parameter values for the maximum conductance 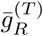, the synaptic weight 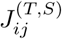, the synaptic reversal potential 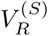, the synaptic decay time constant 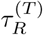, and the amplitudes *A*_1_ and *A*_2_ for the type-2 exponential-decay function in the granular layer, the Purkinje-molecular layer, and the other parts for the CN and IO are shown in Table II.

